# High-accuracy crRNA array assembly strategy for multiplex CRISPR

**DOI:** 10.1101/2024.02.27.582417

**Authors:** Xiangtong Zhao, Lixian Yang, Peng Li, Zijing Cheng, Yongshi Jia, Limin Luo, Aihong Bi, Hanchu Xiong, Haibo Zhang, Jinrui Zhang, Yaodong Zhang

**Author notes:** These authors contributed equally (Xiangtong Zhao, Lixian Yang). Correspondence: Yaodong Zhang, Henan Provincial Key Laboratory of Children’s Genetics and Metabolic Diseases, Children’s Hospital Affiliated to Zhengzhou University, Henan Children’s Hospital, Zhengzhou Children’s Hospital, Zhengzhou, Henan, China Correspondence: Xiangtong Zhao, Henan Provincial Key Laboratory of Children’s Genetics and Metabolic Diseases, Children’s Hospital Affiliated to Zhengzhou University, Henan Children’s Hospital, Zhengzhou Children’s Hospital, Zhengzhou, Henan, China Correspondence: Lixian Yang, Cancer Center, Department of Radiation Oncology, Zhejiang Provincial People’s Hospital (Affiliated People’s Hospital), Hangzhou Medical College, Hangzhou, Zhejiang, China.

## Abstract

Simultaneous targeting of multiple loci with CRISPR system, a tool known as multiplex CRISPR, offers us more feasibility to manipulate and elucidate the intricate and redundant endogenous networks underlying complex cellular functions. Owing to the versatility of the continuously emerging Cas nucleases and the utilization of CRISPR arrays, multiplex CRISPR has been implemented by numerous *in vitro* and *in vivo* studies. However, a streamlined practical strategy for CRISPR array assembly with both convenience and accuracy is still lacking. Here, we present a novel high-accuracy, cost- and time-saving strategy for CRISPR array assembly. Using this strategy, we efficiently accomplished the assembly of up to twelve crRNAs (for AsCas12a) or fifteen crRNAs (for RfxCas13d) in a single reaction. Moreover, we reveal that CRISPR arrays driven by Pol II promoters exhibit a distinct targeting pattern compared to Pol III promoters, which could be exploited for specific distributions of CRISPR intensity. For long CRISPR arrays, we designed a better approach than simply selecting one from U6 or EF1a. Collectively, our study provides a flexible and powerful tool for the convenient implementation of multiplex CRISPR across DNA to RNA, facilitating the dissection of sophisticated cellular networks and the future realization of multi-target gene therapy.

## INTRODUCTION

Over the past decade, following the pioneering discovery and eukaryotic implementation of the CRISPR/Cas9 system ^1–3^, a growing number of new CRISPR systems have been continuously revealed ^4^. And, as the substrate of the best-known Cas9, DNA is no longer the only target within our reach. With the discovery of various RNA-targeting Cas nucleases ^5–11^, RNA can now also be precisely targeted, further extending our range of genetic manipulation from genome to transcriptome. Moreover, by combining with transcriptional activators ^12^, repressors ^13^, base editors ^14^, epigenetic enzymes ^15^, reverse transcriptase ^16^, and tagging molecules ^17^, these RNA-guided effectors have magnificently expanded the molecular toolbox of our basic researchers. Most importantly, CRISPR systems have demonstrated their unlimited potential in the treatment of a growing number of incurable genetic diseases ^18–23^.

As indicated by the full name of CRISPR, the natural CRISPR RNA (crRNA) in bacteria and archaea is usually not transcribed individually, but in a clustered preform from a genomic locus called the CRISPR array, which consists of a succession of direct repeats (DRs) separated by distinct spacers ^4^. Subsequently, the transcribed pre-crRNA is processed into a series of mature crRNAs, each containing a single spacer and a truncated DR. Unlike Cas9, which requires an additional transactivating CRISPR RNA to form a final guide RNA with the crRNA (usually replaced in practice by a chimeric single guide RNA), many other CRISPR systems use the mature crRNAs directly as their guide RNAs . Moreover, the Cas nucleases of these systems typically possess the ability to process their own pre-crRNAs ^4^. These two features render these Cas effectors more amenable to be employed in multiplex CRISPR, as a single array is sufficient to express all the required crRNAs.

Current strategies for assembling customized CRISPR arrays are generally based on the conventional method of cloning a single crRNA, except that the number of annealed oligos is simply multiplied. In fact, such theoretically feasible strategies may work when assembling only a few crRNAs, but are quite incompetent when assembling more, as also demonstrated here. Therefore, an accurate, efficient and practical strategy for CRISPR array assembly is warranted. In the present study, we designed a novel cost-saving strategy for CRISPR array assembly and confirmed its superior accuracy over current alternatives with AsCas12a system. While preserving its accuracy as much as possible, we made several simplifications and optimizations to make this strategy more user-friendly. Next, we employed this strategy in the assembly of crRNAs for an RNA-targeting CRISPR system, highlighting its general applicability and impressive flexibility in addition to high accuracy. When attempting to harness Pol II promoters for CRISPR array expression, we found a distinct targeting pattern of CRISPR arrays transcribed by EF1a (Pol II promoter) compared to the more commonly used U6 (Pol III promoter). For long CRISPR arrays (>200nt), we designed a better approach than simply selecting one from U6 or EF1a. These tandem, hierarchical arrays achieved improved targeting efficiency in CRISPR systems that require abundant expression of crRNAs. Finally, by re-examining the strategy of co-expressing Cas nuclease and CRISPR array on a single transcript, we revealed an unavoidable detrimental effect on the expression of Cas nuclease, despite its unexpectedly satisfactory targeting performance with some CRISPR systems in certain applications.

## RESULTS

### Design of a novel strategy for CRISPR array assembly

Given that it is more convenient and cost-saving when probing the targeting efficiency, we decided to use catalytically dead AsCas12a fused to an artificial VP64-p65-Rta transcription activator (dAsCas12a-VPR, abbreviated as d12a-VPR hereafter) for the following experiments. Besides, an enhanced variant (denAsCas12a-VPR, abbreviated as den12a-VPR hereafter) was also used ^24^. Similar with Cas9-based CRISPR activators ^25–28^, combination of multiple crRNAs enables synergistic activation of endogenous gene targets mediated by d12a-VPR and den12a-VPR (Figure S1). Next, we sought to assess the accuracy of different strategies in the assembly of crRNAs for AsCas12a.

Current strategies for assembly of multiple crRNAs typically consist of two main procedures (Figure S2): annealing of pre-designed single-stranded DNA oligos to form double-stranded DNA (dsDNA) with desired sticky ends, followed by sequential ligation into cloning vectors. Due to their similar principles, we classify them here as Anneal-based strategies.

The accuracy of these types of assemblies relies on the precise sequential ligation of successfully annealed oligos. In practice, however, not all oligos eventually end up annealed to their respective partners as desired, that is, a fair number of oligos remain single-strand after annealing. These single-strand oligos can be a source of trouble, as their natively exposed ends are identical to the sticky ends of the annealed dsDNA, meaning that any site of the sequential ligation can be occupied by them, leading to irreversible premature termination of the assembly. This intrinsic defect is likely to be the dominant cause limiting the number of crRNAs that can be assembled into an array in a single reaction using Anneal-based strategies. Indeed, our exploratory experiments suggested that only up to 6 crRNAs can be efficiently assembled in a single assembly (data not shown).

To push this limit, we designed a novel strategy for CRISPR array assembly that aims to increase both accuracy and maximal number by eliminating perturbations of single-strand oligos (Figure 1). In brief, several additional bases containing BsaI recognition site were appended to the 5’ end of each single-strand oligo, so that the designated inner sticky end was not exposed until a dsDNA was formed and cut by BsaI (Figure S3). These short dsDNA segments can be generated by mainly two alternative methods: (1) Annealing of complete complementary oligo pair, which will be particularly long due to the need for BsaI recognition sites at both ends of the resulting dsDNA; (2) PCR-based approach using much shorter oligo pair that is partially complementary to each other, or to a certain template. We chose the latter because it is cost-saving and also provides us with an opportunity to examine the feasibility of an annealing-free strategy that is sharply different from the conventional Anneal-based methods. To this end, we succinctly referred to our novel approach as PCR-based strategy.

**Figure 1.**
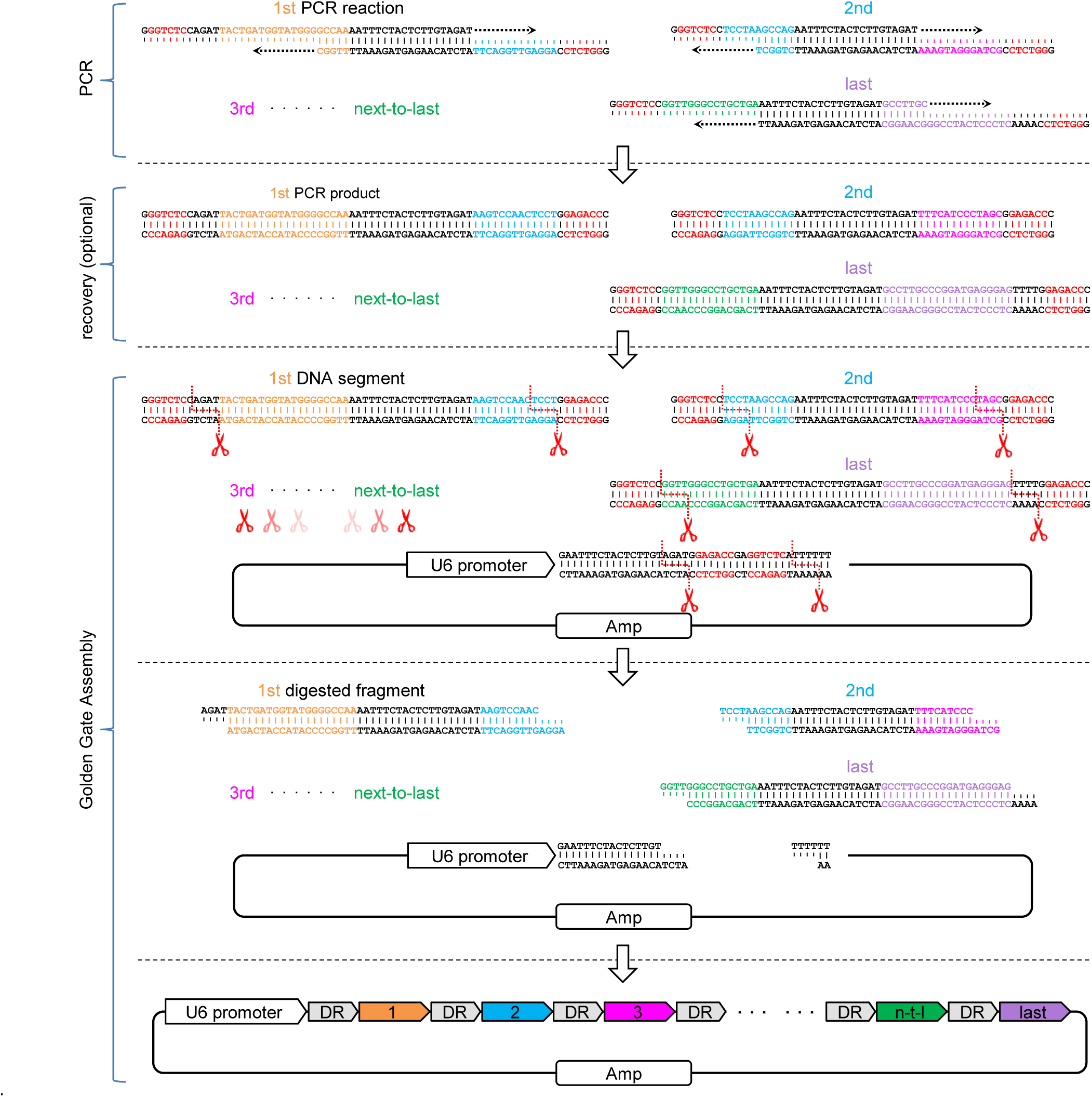
Schematic illustration and workflow of PCR-based CRISPR array assembly strategy. First, set up and run PCR reactions with pre-designed oligo pairs that are partially complementary to each other. Next, an optional recovery step can be performed if necessary (Due to the relatively short DR length of AsCas12a, some of the resulting dsDNA segments were too short to recover using common DNA recovery kits, so we instead used a relatively crude method via ethanol precipitation. Although single-strand oligos or dNTPs might still be included, most of the salts would be removed and the DNA polymerase would be inactivated). Finally, set up and run a standard Gold Gate assembly reaction with purified segments (or diluted PCR products) and destination cloning vector.

No more particular procedures follow the PCR reaction, just optional routine recovery of DNA segments and subsequent standard Golden Gate Assembly.

### High-accuracy assembly of CRISPR array with PCR-based strategy

As for the conventional Anneal-based strategies, there are several versions of the detailed protocol, which differ slightly from each other (with/without pre-digestion before ligation or re-digestion after ligation). A comparison of the accuracy across them was conducted by performing the assembly of 6 crRNAs. While no correct clones were obtained when the ligation mixture was directly transformed into Escherichia coli (E.coli), an additional re-digestion with BsaI between ligation and transformation dramatically increased the accuracy of the selected clones (Figure S4). Moreover, compared to the remaining two, the assembly using standard Golden Gate cycles exhibited improvement or non-inferiority without the procedure of pre-digesting or recovering of cloning vector (Figure S4), hence this specified Anneal-based strategy was used for the following evaluation and comparison.

Next, using our PCR-based strategy, we performed the assembly of an array containing 6 crRNAs with parallel control using Anneal-based strategy for comparison. While all the 60 clones were correct when using PCR-based strategy, the assembly using Anneal-based strategy only achieved a mean accuracy of 37% (P<0.0001) (Figure 2A, Figure S5A). When we increased the number of crRNAs to 7, the accuracy of Anneal-based assembly strategy experienced a sharp decrease, with only one correct out of 60 clones. In contrast, even though a significant reduction was observed, our PCR-based assembly strategy still preserved a mean accuracy of 73% (Figure 2B, Figure S5B).

**Figure 2.**
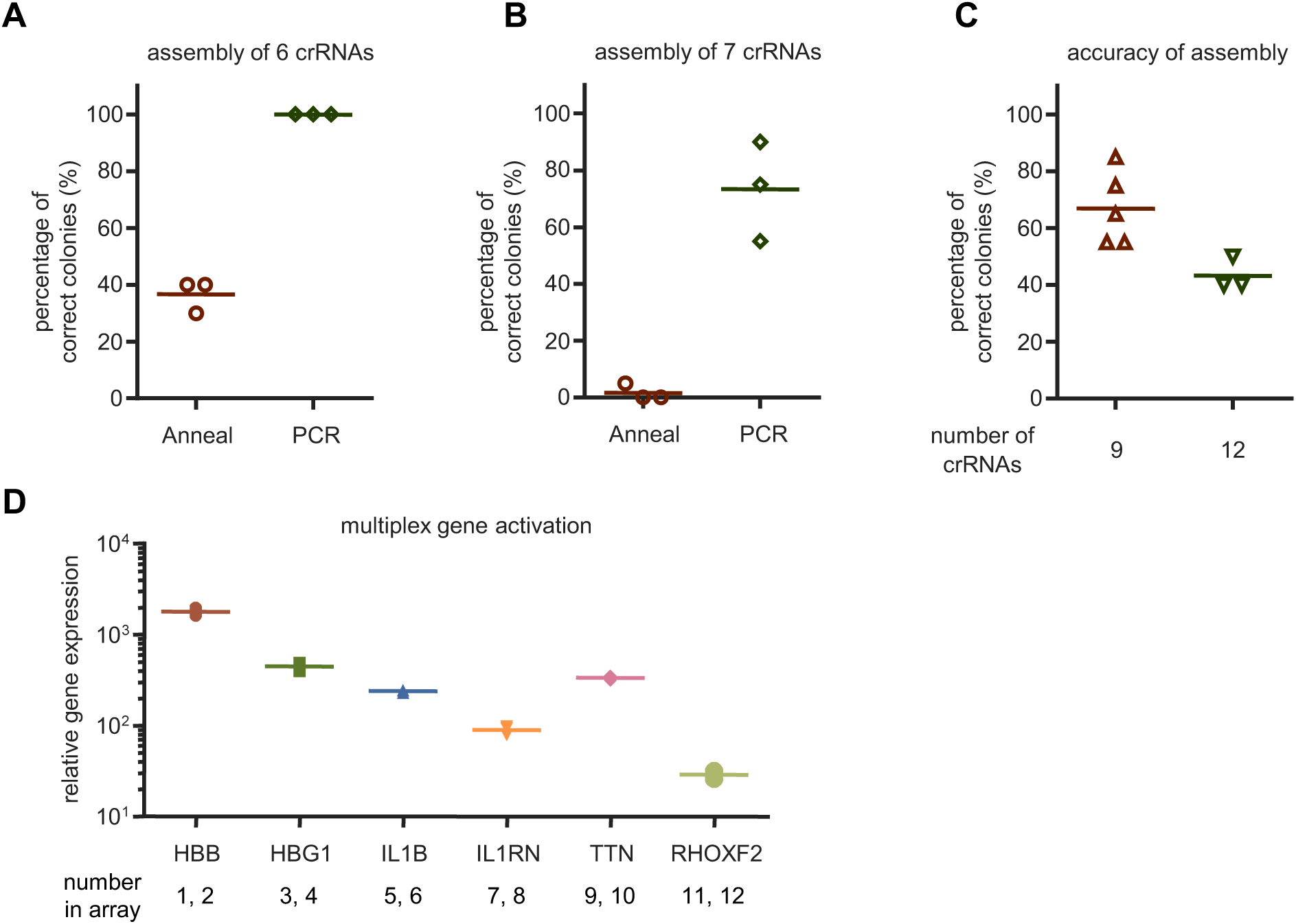
High-accuracy assembly of CRISPR array with PCR-based strategy. (A, B) Accuracies of the assembly of 6 (A) or 7 (B) crRNAs using conventional Anneal-based or novel PCR-based strategy. Values shown as mean, n = 3 independent experiments. (C) Accuracies of the assembly of 9 or 12 crRNAs using PCR-based strategy. Values shown as mean with n ≥ 3. (D) Quantification of relative mRNA expression over non-targeting control in HEK293T cells 48h after transfection with a single plasmid containing CBh-driven denAsCas12a-VPR and U6-driven CRISPR array of 12 crRNAs targeting the indicated 6 genes. CBh, a robust Pol II promoter. Values shown as mean ± sd with n = 3.

Having demonstrated the superiority of our PCR-based strategy over conventional Anneal-based strategies, we then decided to further increase the number of crRNAs to be assembled until the accuracy dropped to a level significantly <50%. Following this criterion, and considering that the extremely low accuracy in assembling 7 crRNAs was already practically meaningless, Anneal-based strategy was no longer evaluated for assembling of more crRNAs. With PCR-based strategy, arrays of 9 or 12 crRNAs were efficiently assembled with mean accuracy of 67% or 43%, respectively (Figure 2C, Figure S5C). Even arrays containing more than 12 crRNAs may still be assembled in a single reaction, but in view of the gradual decrease in assembly accuracy as the number of crRNAs increases, the practical value of such experiments would be limited, thus we ceased here and determined the maximum number of crRNAs for AsCas12a that could be assembled into an array in a single reaction to be 12.

The resulting array of 12 crRNAs contained spacers targeting promoters of 6 endogenous human genes (2 crRNAs for each gene): *HBB*, *HBG1*, *IL1B*, *IL1RN*, *TTN*, *RHOXF2*. Together with this array, den12a-VPR achieved robust activation of these six genes simultaneously in HEK293T, with a range of ∼20 to ∼2000 fold (Figure 2D).

### Simplification and optimization of the PCR-based strategy for CRISPR array assembly

While our PCR-based assembly strategy exhibited greater accuracy than other alternatives, the requirement of an additional recovery procedure of PCR products may hinder its wider application. To overcome this inconvenience, we next sought to circumvent this procedure while preserving the accuracy of the assembly as much as possible. Instead of the time-consuming recovery to obtain purified dsDNA segments in the standard assembly protocol, PCR products were diluted and then added directly to the Golden Gate assembly reaction. From a wide range of PCR primer concentrations and volumes of PCR products pipetted for subsequent assembly, the best accuracy was achieved when using a final primer concentration of 2.5uM for PCR and 0.1uL (i.e., 1uL after 10-fold dilution) from each of the resulting PCR mixtures for a 20uL assembly reaction (for the designated DNA polymerase we used, data not shown). Assemblies of 6, 7, 9, and 12 crRNAs were performed following this reaction condition. The mixing of unwanted PCR components other than dsDNAs into the subsequent assembly reaction did compromise the final accuracy, but only to a modest degree, with the exception of the assembly of 12 crRNAs, from which correct clones were barely obtained (Figure S6). Nevertheless, this simplified recovery-free version of our PCR-based strategy still outperformed current alternative assembly strategies without the requirement for additional procedures, and can be a rational choice when assembling no more than 9 crRNAs.

Another shortcoming, which may make our assembly strategy complicated and confusing, comes from the DR sequence of AsCas12a. The 19nt DR of AsCas12a has a relatively low melting temperature (44∼45°C, Tm) due to its low GC content (∼26%). This intrinsic property makes it somewhat challenging to design initial primers for subsequent PCR reactions, since oligos must be extended beyond the DR sequence to obtain a minimal appropriate Tm (50∼55°C). To address this issue, we next sought to introduce mutations into the original wild-type DR, with the aim of increasing its Tm value to a proper extent (>50°C) without compromising its function. Indeed, several previous studies have made such attempts, but with distinct aims ^29–31^. Based on their findings, we generated 3 variants of wild-type DR (Figure 3A). Among these candidates, one variant with a Tm of 49°C showed non-inferiority in targeting efficiency compared to the wild-type DR (Figure 3B). Although the Tm of this variant is still slightly below 50°C, it is already feasible to use the DR sequence directly as the end of primers and 45°C as the annealing temperature for subsequent PCR reactions, thus simplifying the primer design procedure (Figure 3C). Based on this DR variant, arrays of 9 and 12 crRNAs were successfully assembled with mean accuracy of 75% and 15%, respectively (Figure 3D, Figure S7). Moreover, the assembly of an array containing 9 crRNAs was also accomplished efficiently when both mutant DR and the recovery-free version of PCR-based strategy were employed (Figure S8).

**Figure 3.**
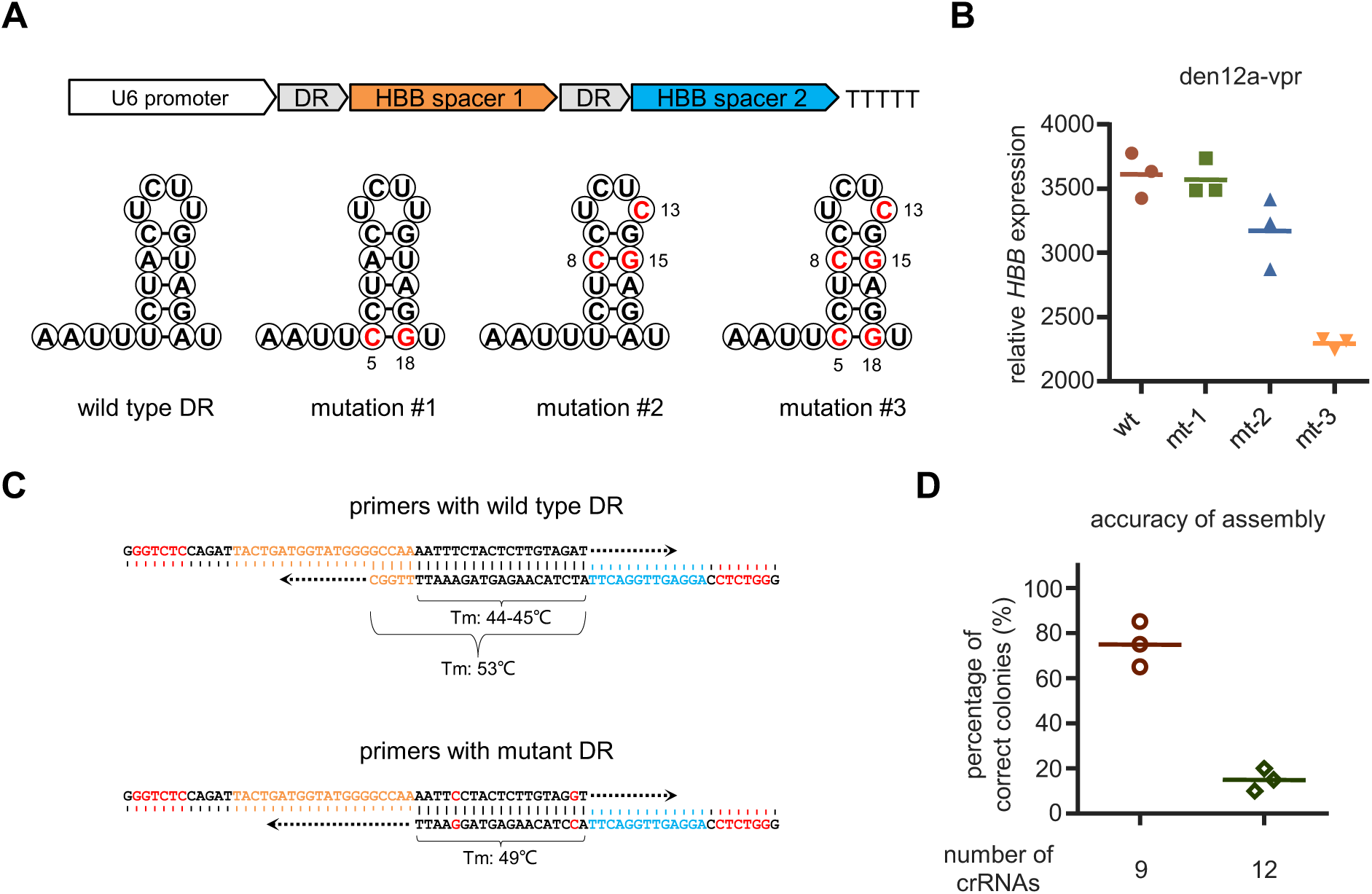
Simplification of the PCR-based strategy for CRISPR array assembly. (A) Schematics of wild type and 3 mutant DR variants of AsCas12a system. (B) Quantification of relative *HBB* expression over non-targeting control in HEK293T cells 48h after transfection with den12a-VPR and arrays with DR variants depicted in (A). (C) An illustrative example showcasing the streamlining of the initial primer design through the utilization of mutant DR with higher GC content. (D) Accuracies of the assembly of 9 or 12 crRNAs with mutant DR using PCR-based strategy. Values shown as mean with n = 3.

### High-accuracy assembly of CRISPR array for a Cas13d nuclease

While the previous content has centered around AsCas12a, the potential applications of our assembly strategy can be extended far beyond this specified CRISPR system. Theoretically, owing to its outstanding flexibility and generalizability, this crRNA array assembly strategy can be applied to most CRISPR systems whose crRNA contains a DR followed by a spacer, especially when the Cas nuclease has the ability to process its own crRNA.

To confirm this concept, we next sought to employ it to assemble crRNAs of another distinct CRISPR system, Cas13d, a family of RNA-targeting Cas nucleases ^4^. Among them, RfxCas13d was chosen for the subsequent assessment ^9^. The commonly used crRNA for RfxCas13d is composed of a 36nt DR and a spacer ranging from 22 to 30nt. We selected an intermediate spacer length of 26nt for the following CRISPR array assembly. Distinct from that of AsCas12a, DR of RfxCas13d is much longer, and has a modest GC content (50%). These properties render the assembly of its crRNAs more suitable for our strategy: (1) Design of the primers will be much easier than that of AsCas12a, as all the forward and reverse primers can use the same ∼20 bases 3’ section, respectively; (2) Using current alternatives for such a CRISPR system with long DR sequence would be considerably more costly, since it would be unavoidable to purchase particularly long oligos (>60 bases), which are usually several times more expensive per base than the shorter routine oligos due to their complex production process. The detailed procedure is generally the same as the assembly of crRNAs for AsCas12a, with two minor modifications (Figure S9). The resulting procedure is somewhat reminiscent of several previous strategies for assembling sgRNA cassettes used in Cas9-based multiplex CRISPR ^32–34^.

Surprisingly, our PCR-based assembly strategy performed better with RfxCas13d compared to AsCas12a, as an array of up to 15 crRNAs could be assembled in one pot with an acceptable accuracy (Figure 4A, Figure S10). This improved performance might be owing to purer dsDNA segments or other uncertain reasons. With this array, we attempted to achieve simultaneous cleavage of 15 endogenous transcripts. 48h after transfection, most of these transcripts were efficiently cleaved detected with corresponding primers flanking the cleavage sites (Figure 4B).

**Figure 4.**
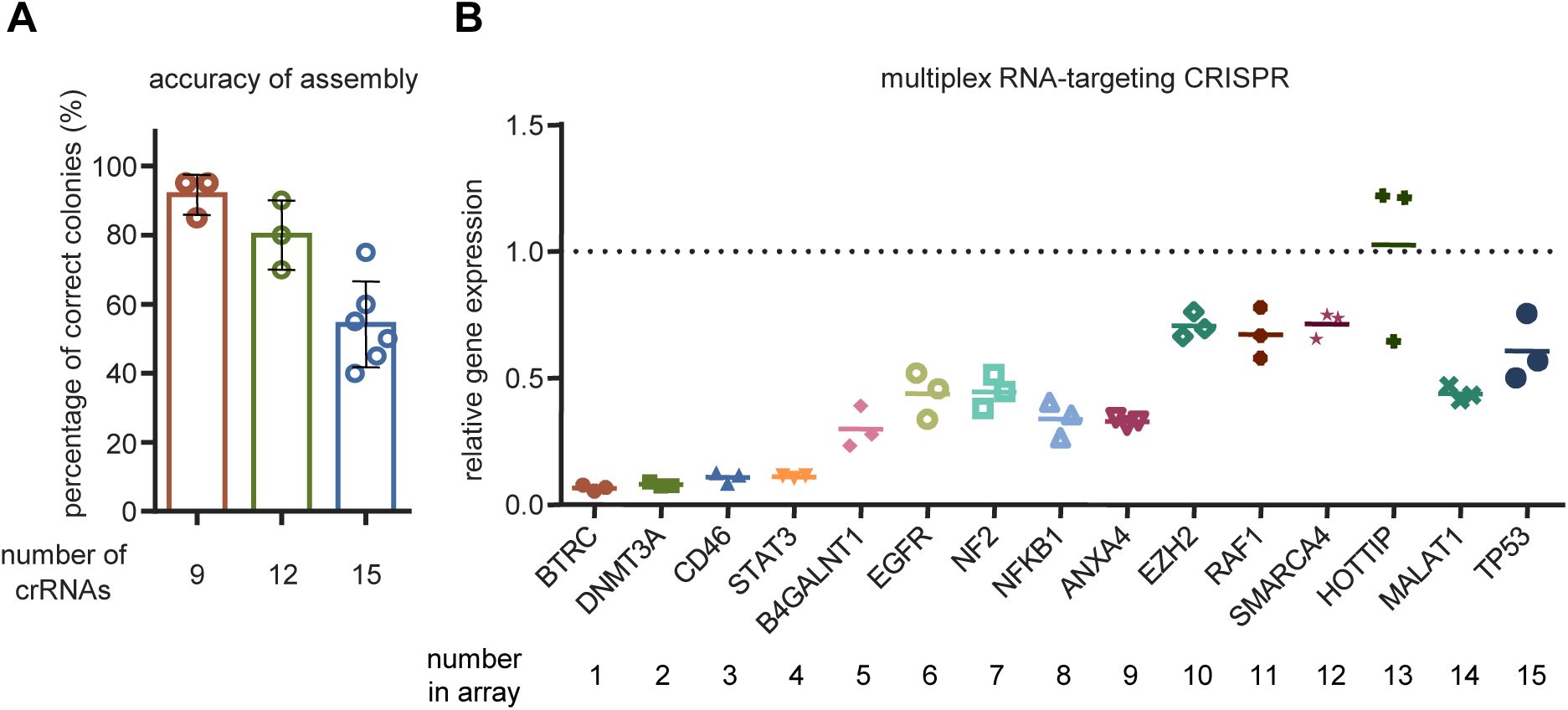
High-accuracy assembly of crRNAs for a Cas13d nuclease. (A) Accuracies of the assembly of 9, 12, or 15 crRNAs for RfxCas13d using PCR-based strategy. Values shown as mean with n ≥ 3. (B) Quantification of relative mRNA expression compared to non-targeting control in HEK293T cells 48h after transfection with a plasmid containing EF1a-RfxCas13d and U6-driven CRISPR array of 15 crRNAs targeting the indicated 15 endogenous transcripts. Values shown as mean with n = 3.

### Distinct targeting patterns of Pol II/III promoter-driven CRISPR arrays

Although the array of 15 crRNAs used for RfxCas13d had reached a length of ∼1200nt, the U6 promoter still seemed capable of driving its transcription. This may be a bit discrepant with our conventional conception that Pol III promoters (for example, U6, H1, 7SK) are typically employed for the transcription of small RNAs, although RNA up to 800 nucleotides has also been reported to be efficiently transcribed by U6 ^35^. One solution to this indeterminate restriction is to replace Pol III promoters with Pol II promoters (for example, CMV, EF1a), since they are capable of transcribing much longer RNAs. Several reports have conducted the evaluation of harnessing Pol II promoters to express crRNAs, or synthetic sgRNAs ^36, 37^.

To determine whether CRISPR arrays transcribed by Pol II promoters work as efficiently as those by Pol III promoters, we transfected HEK293T with den12a-VPR and an array of 12 crRNAs driven by either U6 (U6-array) or EF1a (EF1a-array), followed by quantification of the expression of the corresponding 6 targets 48h post-transfection. Unexpectedly, no straightforward determination of superiority or inferiority could be drawn from the results, as each had its own strengths and weaknesses (Figure 5A). Nonetheless, if we had to make a conclusion, it seemed that U6 favored the expression of the first few crRNAs. When targeting individual genes using 2 crRNAs each, U6 outperformed EF1a for both *HBB* and *RHOXF2* (Figure 5B and C), suggesting the superiority of U6 over EF1a when driving the expression of short CRISPR arrays. Moreover, since we could not rule out the possibility that crRNAs expressed from U6 were already saturated, the disparity observed here might been underestimated. Indeed, a greater gap could been seen by varying the ratio of den12a-VPR to the array (Figure S11). However, the disparity between these promoters was not embodied in gene editing using nuclease-active AsCas12a (Figure 5D). The reason might be that the number of targets in the genome is limited, and once edited with indel, CRISPR elements were no longer needed, meaning that a small amount of crRNAs was already sufficient and more would be redundant.

**Figure 5.**
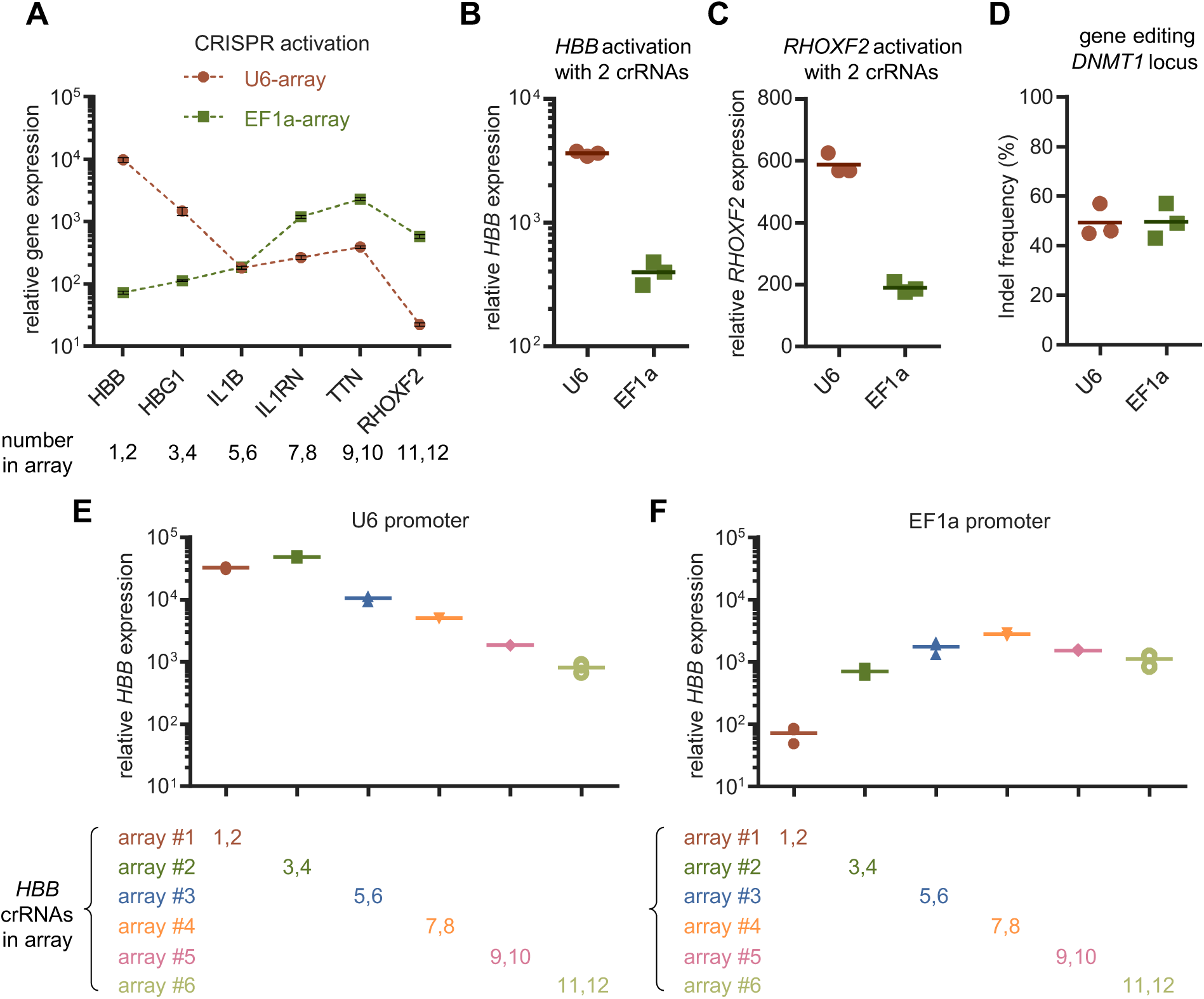
Distinct targeting patterns of Pol II/III promoter-driven CRISPR arrays. (A) Quantification of relative mRNA expression over non-targeting control in HEK293T cells 48h after transfection with EF1a-denAsCas12a-VPR and an array of 12 crRNAs driven by either U6 (U6-array) or EF1a (EF1a-array). Values shown as mean ± sd with n = 3. (B, C) Quantification of the indicated gene expression over non-targeting control in HEK293T cells 48h after transfection with EF1a-denAsCas12a-VPR and U6- or EF1a-driven array of 2 crRNAs targeting the promoter of *HBB* (B) or *RHOXF2* (C). (D) Quantification of gene editing efficiency in HEK293T cells 72h after transfection with enAsCas12a and a *DNMT1*-targeting crRNA driven by either U6 or EF1a. (E, F) Quantification of relative *HBB* expression over non-targeting control in HEK293T cells 48h after transfection with EF1a-denAsCas12a-VPR and the indicated arrays (12×) driven by U6 (E) or EF1a (F), where HBB-targeting crRNAs were positioned at different loci. Values shown as mean with n = 3.

To determine whether this phenomenon represented a general pattern, we continued the comparison with other CRISPR systems. 48h after transfection with RfxCas13d and an array of 15 crRNAs driven by either U6 or EF1a, no conclusions could be drawn except that U6-array showed superiority in the targeting of the first 4 transcripts (Figure S12A). When harnessed for the expression of single crRNA, EF1a exhibited a significantly lower target cleavage efficiency compared to U6 (Figure S12B).

In contrast to EF1a-array, the location of crRNAs in the U6-array appeared to have a greater influence on their own expression. When the internal order of the CRISPR array was thoroughly reversed, U6-array experienced a more drastic change in its multiplex CRISPR efficiency (Figure S13). To obtain an intuitive and precise correlation between transcriptional strength and distance from the promoter, the two crRNAs targeting HBB promoter were placed at a series of different loci, while the overall component was kept constant (Figure 5E and F). The first few crRNAs of U6-array achieved a very high level of expression, a level that could not be reached by EF1a. Thereafter, the intensity of the transcription dropped off continuously with increasing distance from U6. On the contrary, distance from EF1a promoter had a much weaker effect on the expression of crRNAs, except that the first few crRNAs might undergo inadequate expression.

These results suggest that Pol II and III promoters are not functionally equivalent to each other when driving the expression of CRISPR arrays. Specifically, while U6 favors the expression and execution of the first few crRNAs within ∼200nt, EF1a seems to distribute its relatively milder strength more evenly. For single crRNA or small CRISPR arrays (<200nt), especially when using the RfxCas13d system, harnessing Pol II promoters may still be an inadequate alternative unless its mild expression is acceptable.

### Improve the targeting efficiency by manipulating the architecture of long CRISPR arrays

Having revealed the superiority of U6 over EF1a when transcribing the first few crRNAs within ∼200nt, we next decided to find a better approach for the transcription of longer CRISPR arrays than simply selecting one from U6 or EF1a. By splitting long CRISPR array into several short ones, each of them could be assigned with a U6 promoter, thus employing the edge of U6 in the strength of transcribing short RNAs, while circumventing its disadvantage in transcribing long RNAs. The principle and workflow is similar to the assemblies previously performed in this study, with the only difference that small PCR segments containing crRNAs and longer PCR segments containing U6 will be mixed and assembled in one reaction (Figure S14). However, we failed to assemble the correct array matching our expectations on our maiden attempt (data not shown). This might be attributed to the introduction of repeated U6 elements, which made the resulting array more prone to go through recombination events.

In previous content, sometimes we amplified the array of 12 or 15 crRNAs as an intact sequence from purified plasmid that already contained the entire array (considered as the second round of PCR), and then cloned this unique insert into a new vector (considered as the second round of assembly), a simple procedure to transfer an existing array from one vector to another.

Although the accuracy of assembling 12 or 15 individual crRNAs into one array in the first round could be as low as 15% (Figure 3D), the second round of assembly always achieved an accuracy of >80% (data not shown). One primary determinant of this disparity may be that as the number of inserts increases during Golden Gate Assembly, it becomes more and more challenging for the DNA ligase to seal each gap; following the transformation of assembly mixture into competent cells, plasmids with gap(s) will induce the DNA repair function in E.coli, a progression that often leads to recombination. Thus, reducing the number of inserts may be one solution to frequent recombination events. To test this hypothesis, we decided to conduct the second round of assembly. For the sake of time, assembly mixture of the first round was directly used as the PCR template of the second round, instead of validated and purified plasmid (Figure S15). The mean accuracy of the assembly of the array with 12 crRNAs was improved to 88% (Figure S16). However, we must point out that amplifying the entire array from the crude mixture of previous assembly may not always succeed, especially when the destination fragment spans more than a dozen crRNAs. An array with 20 crRNAs was successfully assembled in two rounds, but we had to perform two individual amplifications with two pairs of primers in the second round (Figure S16).

With the concept of improving the accuracy by performing an extra round of assembly, we decided to confront the obstacle of assembling hierarchical CRISPR arrays again. In the second round, RfxCas13d CRISPR array consisting of four U6-crRNAs units was successfully assembled with mean accuracy of 33% (Figure S17B), which had been shown to be impossible in the first round. Similarly, this strategy significantly increased the accuracy of the assembly of denAsCas12a CRISPR array consisting of three U6-crRNAs units (Figure S17A).

In contrast to arrays driven by a single U6 or EF1a, a tandem array driven by three U6 achieved robust transcriptional activation for all of the six targets in denAsCas12a-VPR mediated multiplex CRISPR system (Figure 6A and B). Similarly, for most of the fifteen endogenous transcripts, RfxCas13d showed a more prominent cleavage efficiency when working with a tandem array driven by four U6 (Figure 6C and D).

**Figure 6.**
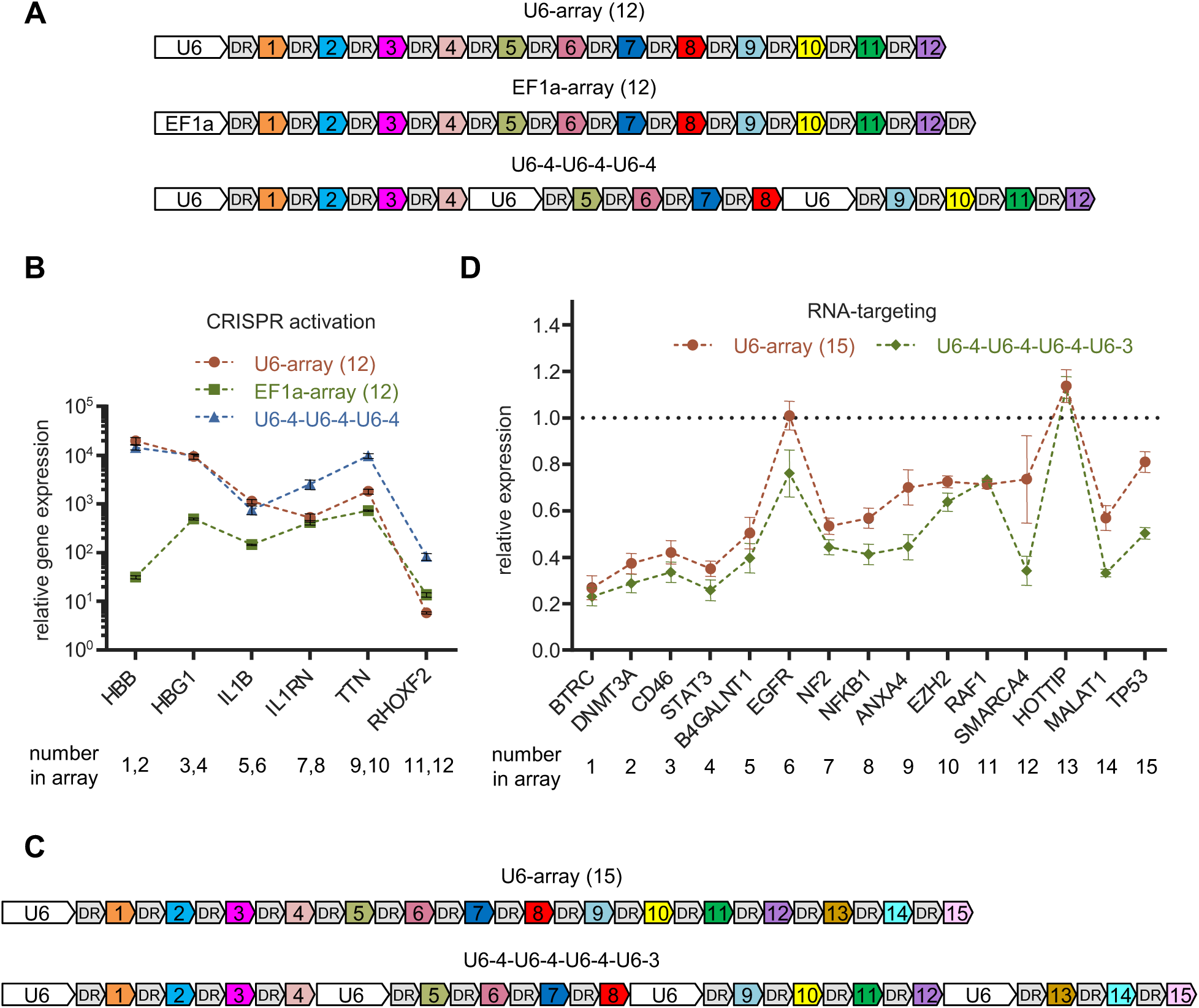
Improve the targeting efficiency by optimizing the internal architecture of the CRISPR array. (A) Schematics of CRISPR arrays used with denAsCas12a-VPR: U6-array (12), EF1a-array (12), U6-4-U6-4-U6-4. (B) Quantification of relative mRNA expression over non-targeting control in HEK293T cells 48h after transfection with EF1a-denAsCas12a-VPR and the indicated arrays. Values shown as mean ± sd with n = 3. (C) Schematics of CRISPR arrays used with RfxCas13d: U6-array (15), U6-4-U6-4-U6-4-U6-3. (D) Quantification of relative mRNA expression of the indicated genes compared to non-targeting control in HEK293T cells 48h after transfection with RfxCas13d and the indicated arrays. Values shown as mean ± sd with n = 3.

### Co-expression of Cas nuclease and CRISPR array on a single transcript

Now that Pol II promoters are capable of driving the expression of CRISPR arrays in addition to protein-encoding genes, one might consider the possibility of co-expressing Cas protein and CRISPR array on a single transcript. To achieve such kind of co-expression, the coding sequence of Cas protein must be positioned upstream of the CRISPR array, followed by a poly(A) tail at the end. However, this arrangement poses a problem: processing of the CRISPR array will inevitably leads to the separation of Cas protein mRNA from the poly(A) tail, ultimately resulting in inadequate expression of Cas protein and unsatisfactory CRISPR efficiency. In fact, a previous study has attempted to address this issue by harnessing a putative mRNA-stabilizing element called “Triplex” ^37^. Their results showed that the introduction of “Triplex” between EGFP and CRISPR array completely rescued the loss of EGFP fluorescence due to the processing of downstream array. Frankly, we have to express a bit of concern about this kind of “complete rescue”, as it challenges the irreplaceable role of poly(A) tails in stabilizing eukaryotic mRNA ^38^.

To test these suspicions, we decided to further investigate whether this specific co-expression pattern is a viable option for multiplex CRISPR. Consistent with conventional understanding, removal of the poly(A) tail remarkably reduced EGFP expression at both mRNA and protein levels (Figure S18A and B), emphasizing the crucial role of poly(A) tail in stabilizing mRNA. To examine the mRNA-stabilizing efficacy of Triplex and the well-known WPRE element, we cloned EGFP downstream of an EF1a promoter, followed by an array of 12 crRNAs for AsCas12a, with or without a Triplex or WPRE element between EGFP and CRISPR array. The resulting plasmids were transfected into HEK293T together with control vector or den12a-VPR. 48h post-transfection, as a result of the processing of crRNAs, simultaneously expressed den12a-VPR dramatically reduced EGFP mRNA and fluorescence. Distinct from the previous study ^37^, additional introduction of Triplex or WPRE resulted in mild, if any, improvement in the mRNA or fluorescence of EGFP (Figure S18C and D).

Next, we decided to directly evaluate the targeting efficiency of co-expressing den12a-VPR and CRISPR array on a single transcript (Figure S19A). Co-expression via direct integration did result in a significant reduction in den12a-VPR mRNA/protein expression, but not in the ultimate CRISPRa efficiency, which increased instead (Figure S19B-D). This discrepancy between decreased den12a-VPR expression and enhanced CRISPRa efficiency, might be a result of increased expression of CRISPR array due to the insertion of upstream den12a-VPR. Indeed, when EF1a-array and den12a-VPR were delivered by individual constructs, the expression of CRISPR array and the final CRISPRa efficiency could be enhanced by inserting an EGFP-coding sequence upstream of CRISPR array, but not by inserting a random stuffer (Figure S20A). This phenomenon is not unique, as CMV-driven CRISPR array could also be enhanced via this approach (Figure S20B). Moreover, enhanced CRISPRa efficiency could be achieved by increasing the ratio of array-coding plasmid to that expressing den12a-VPR (Figure S21A). To our surprise, even the original version(i.e. den12a-VPR), which is supposed to possess considerably lower activity and require a smaller amount of crRNAs, showed improved CRISPRa efficiency when the ratio of EF1a-array was increased or combined with EF1a-GFP-array (Figure S21B).

As for additional insertion of Triplex or WPRE when co-expressing den12a-VPR and CRISPR array on a single transcript, while WPRE seemed to slightly rescue den12a-VPR mRNA/protein, Triplex did not (Figure S19B and C). However, neither of them improved the CRISPRa efficiency compared to direct integration (Figure S19D).

Taken together, the strategy of co-expressing Cas protein and CRISPR array on a single transcript may be a viable option for some CRISPR systems under certain circumstances, but not always. An important determinant, according to our results, is that the optimal ratio of Cas effector to its crRNA varies across different CRISPR systems. Systematic and rigorous evaluation is needed before applying this particular approach to other CRISPR systems not tested here, especially for *in vivo* delivery, which is lacking in our study.

## DISCUSSION

Although up to 15 crRNAs could be assembled in a single reaction using our method, some previous studies did use arrays containing much more crRNAs, even>20 ^37^. However, they typically performed more than one round of assembly or were relied on high-cost long dsDNA segments purchased from companies. Based on purified construct coding the array of 12 crRNAs assembled in the first round (while the assembly product of the first round was used as template in Figure S15), we performed a second round of PCR and subsequent assembly (Figure S22A). Although evident recombination events occurred during clonal expansion since we used the same template for all three PCR reactions, the array of 36 crRNAs was efficiently assembled with high accuracy (Figure S22B and C). Due to a lack of appropriate application examples, we did not further determine the maximum number of crRNAs that could be assembled in two rounds of assembly. Even so, given the remarkable accuracy in assembling 36 crRNAs, the successful assembly of more crRNAs (i.e., 40, 50, or even more) within two rounds is quite possible.

Speaking of unwanted recombination events during the transformation of plasmids into competent E.coli, we ourselves are still puzzled by its “elusiveness” to this day. For convenience, even for the transformation of lentiviral constructs, we routinely use DH5α, a commonly used recA1 mutant competent E.coli strain, instead of stbl3 or NEB stable, which are supposed to further reduce recombination events between repeat elements in the plasmid. A lower temperature of 30℃ for bacterial culture was also rarely used. But the fact is that we have never encountered any recombination events between the two LTRs (long terminal repeat) of lentiviral constructs. Specific to this study, the accuracy of the assembly of crRNAs was not improved by using stbl3 or NEB stable, nor by culturing at 30℃ (data not shown). Compared to the transformation of newly assembled mixture, recombination events were quite rare during the re-transformation of repeat-containing plasmids extracted from bacteria (data not shown). Frankly, we are not sure whether most of the negative colonies are the result of unwanted recombination or simply failed assembly. As these reagents associated with Golden Gate cloning have been continuously upgraded over the years ^39, 40^, the limits on the maximum number of crRNAs that can be assembled in a single reaction, if the failure of assembly is currently the primary constraint, may be pushed further in the future.

Distinct from previous report ^24, 37^, our results indicate that EF1a is not functionally equivalent to U6 in the transcription of CRISPR arrays or single crRNA. Despite the difference between their strength when transcribing transcripts of varying length, additional modifications (for example, 7-methylguanosine (m7G) cap, poly(A) tail) may also have an effect on the subcellular localization or processing of the CRISPR array, thereby affecting its final execution.

While the loss of poly(A) tail by different approaches all led to reduced EGFP expression, we noticed that the reduction seemed to be more drastic when the loss of poly(A) tail was caused by the processing of downstream CRISPR array, compared to simply removing SV40-poly(A) signal in the construct (Figure S18). Although we are not sure that the former accurately represents the real effect of poly(A) deficiency, the latter may underestimate it. When SV40-poly(A) is removed from the construct, Pol II promoters do not stop transcribing upon completion of EGFP transcription, but instead continue to transcribe downstream uncertain sequences until terminated for other reasons. Thus, the 3’ terminus of the final EGFP mRNA is not directly exposed, but is “protected” by downstream RNA of uncertain length, which may delay the degradation of EGFP mRNA by acting as a stuffer. This reminds us of lentiviral constructs in which the poly(A) signal between the two LTRs has to be removed as it would lead to premature termination of the viral RNA during virus packaging. Consequently, the sequence integrated into the host genome after lentivirus infection does not have a poly(A) signal. A WPRE element is frequently introduced upstream of the 3’ LTR, which is supposed to stabilize and facilitate the translation of mRNA ^41^. While the improvement in the expression of upstream coding genes in our previous results with transient expression was mild, WPRE markedly enhanced the expression of upstream EGFP in the context of lentiviral delivery, consistent with the conventional concept. Triplex, by contrast, failed under this circumstance (Figure S23).

Although the previously reported co-expression strategy inevitably resulted in reduced expression of Cas effector in our re-evaluation, it did not necessarily lead to compromised CRISPR efficiency, at least in denAsCas12a-VPR mediated multiplex activation system. However, as for the genomic DNA targeting system with the primordial nuclease active AsCas12a (or enAsCas12a), it is currently undetermined whether this particular co-expression approach is compatible.

One of the shortcomings of this study, which we must emphasize, is that all the evaluations were performed *in vitro* using cell lines, and these results may not precisely represent the *in vivo* performance of the CRISPR systems and expression methods assessed here. Therefore, rigorous *in vivo* evaluation is still warranted before further application, especially when harnessing Pol II promoters to express CRISPR array or using the co-expression strategy. Nevertheless, our findings will significantly simplify the preparatory work prior to *in vivo* manipulation of multiple targets and boost the exploration of potential applications using multiplex CRISPR.

## MATERIALS AND METHODS

### Plasmid construction

The coding sequences of AsCas12a, VPR, RfxCas13d were amplified from pY108 (Addgene #84739), pXR001 (#84739), lenti-EF1a-dCas9-VPR-Puro (#99373) respectively. Amplified fragments were cloned into destination vectors using standard digestion-ligation or Gibson methods. Gibson cloning was used to introduce desired mutations into protein-coding plasmids. CRISPR arrays were assembled using direct ligation or standard Golden Gate method. The junction sets of the 4-base overhangs were decided based on their ligation fidelity predicted by the online tool: NEBridge Ligase Fidelity Viewer (https://ggtools.neb.com/viewset/run.cgi) ^39^. Detailed protocols are provided in Supplementary Methods. All constructs were verified by Sanger sequencing. Detailed sequences of these constructs are listed in Supplementary Table.

### Cell culture and transient transfection

HEK293T cells were cultured in Dulbecco’s modified Eagle’s medium (DMEM) supplemented with 10% FBS and 1% penicillin/streptomycin at 37℃ with 5% CO2. For transient transfection, 2×10^5^ HEK293T cells per well were seeded in 24-well plates. When the confluency reached 60%–80% on the following day, a total of 500 ng plasmids were transfected into each well using Lipo8000^™^ Transfection Reagent (Beyotime) according to the manufacturer’s protocol. Transfected cells were supplied with fresh complete DMEM (containing FBS) every 24h, and harvested 48-72h after transfection for downstream experiments.

### Lentivirus production and infection

For lentivirus packaging, 5×10^5^ HEK293T cells per well were seeded in 12-well plates. When the confluency reached 80% on the following day, 1 μg of mixed plasmids (transfer: psPAX2: pMD2.G = 4: 3: 1) were transfected into each well using Lipo8000^™^ (Beyotime). 12-24h after transfection, media was replaced with 1mL fresh complete DMEM. 48h after transfection, another 1mL complete DMEM was supplemented. 72h after transfection, lentivirus-containing supernatant was harvested, then filtered through a 0.45 μm filter (or centrifuged at 2,000 g for 5min) and stored at 4°C (or -80°C for long-term preservation).

For lentivirus infection, 1.5×10^5^ HEK293T cells per well were seeded in 24-well plates. When the confluency reached 40%–60% on the following day, media was replaced with 0.5mL fresh complete DMEM and 1.5mL lentivirus-containing supernatant. 12-24h after infection, media was replaced with fresh complete DMEM, and refreshed/supplemented every 24h until cells were harvested 72h after infection.

### Extraction of RNA and RT-qPCR

Total RNA was extracted using RNAiso plus (Takara) 48-72h after transient transfection or lentivirus infection. 1 μg total RNA was then reverse transcribed using ReverTra Ace^®^ qPCR RT Kit (Toyobo) with supplied primer mix or gene-specific primers (when designated, and sequences are listed in Supplementary Table 2), followed by qPCR using ChamQ Universal SYBR qPCR Master Mix (Vazyme) and primers listed in Supplementary Table 1. qPCR reactions were carried out on LightCycler 480 II (Roche). Quantification of RNA expression was normalized to ACTB (unless otherwise specified) and calculated using the ΔΔCt method.

### Electrophoresis of oligonucleotides

Electrophoresis of DNA oligos was run using 20% polyacrylamide gel in 1×TBE buffer. The gel was then stained with Gel-Red (Beyotime) for 1 h at room temperature, and scanned with a UV transilluminator after washes.

### Quantification of gene editing

72h after transfection, genomic DNA was extracted using Rapid Animal Genomic DNA Isolation Kit (Sangon Biotech) following the manufacturer’s protocol. Extracted DNA was used as template to amplify the target region flanking the edited site with specific primers. PCR amplicons were purified with AxyPrep PCR Clean-up Kit (Axygen) following the manufacturer’s protocol. Purified DNA was then subjected to sanger sequencing using the forward PCR primer as sequencing primer. The resulting “.ab1” files were uploaded to obtain the final quantitative spectrum of the indels using the online tool: https://ice.synthego.com ^42^.

### Western blot

72h after transient transfection or lentivirus infection, cells were lysed in RIPA buffer (MedChem Express) supplemented with PMSF and BeyoZonase^™^ Super Nuclease (Beyotime). Total protein concentration was quantitated using Pierce^™^ BCA Protein Assay Kit (ThermoFisher Scientific). All lysates were diluted to a concentration of 1250 ng/μL, then mixed with 5×loading buffer (a final protein concentration of 1μg/μL) and boiled for 5 min. Equal amounts of protein were loaded and separated by SDS-PAGE, then transferred to PVDF membrane. Membrane was blocked with 5% nonfat milk in TBST buffer for 1 h at room temperature, and incubated with appropriate dilutions of primary antibody overnight at 4°C. The next day, the membrane was washed three times with TBST, then incubated with horseradish peroxidase-conjugated secondary antibody for 1 h at room temperature. After three washes, chemiluminescent signal in membrane was captured using a CCD camera-based imager.

### Statistical analysis

Values are reported as mean or mean ±SD as indicated in the appropriate figure legends. Unless otherwise noted, at least three biological replicates were performed for each experiment. When comparing two groups, the statistical difference was determined by unpaired Student’s t test. One-way ANOVA was used to assess the significance between more than two groups. Two-way ANOVA was used when comparing across two factors. P < 0.05 was considered statistically significant. Prism 6.01 was used for all statistical analyses.

## DATA AVAILABILITY

The authors declare that data supporting the results of this study are available in the article and its supplementary materials. Further resource information is available from the corresponding author upon reasonable request.

## SUPPLEMENTARY INFORMATION

Supplemental information can be found online at.

## ACKNOWLEDGEMENTS

We thank members of the institute of immunology (Zhejiang University School of Medicine) for supporting this research. This work was financially supported by Postdoctoral Starting Grants for Scientific Research from Zhejiang Provincial People’s Hospital (C-2023-BSH28), National Natural Science Foundation of China (82102814) and Zhejiang Provincial Natural Science Foundation of China (LQ22H160053).

## AUTHORS CONTRIBUTIONS

XZ: conception and design, collection of data, data analysis, manuscript writing; LY: conception and design, collection of data, data analysis, manuscript writing, financial support; PL, ZC, YJ, LL, AB, HX, HZ, JZ: collection of data, data analysis, manuscript editing, administrative, technical, material or financial support; YZ: conception and design, data analysis, financial support, revision and final approval of manuscript.

## DECLARATION OF INTERESTS

The authors declare no competing interests.

---

**graphical abstract.**
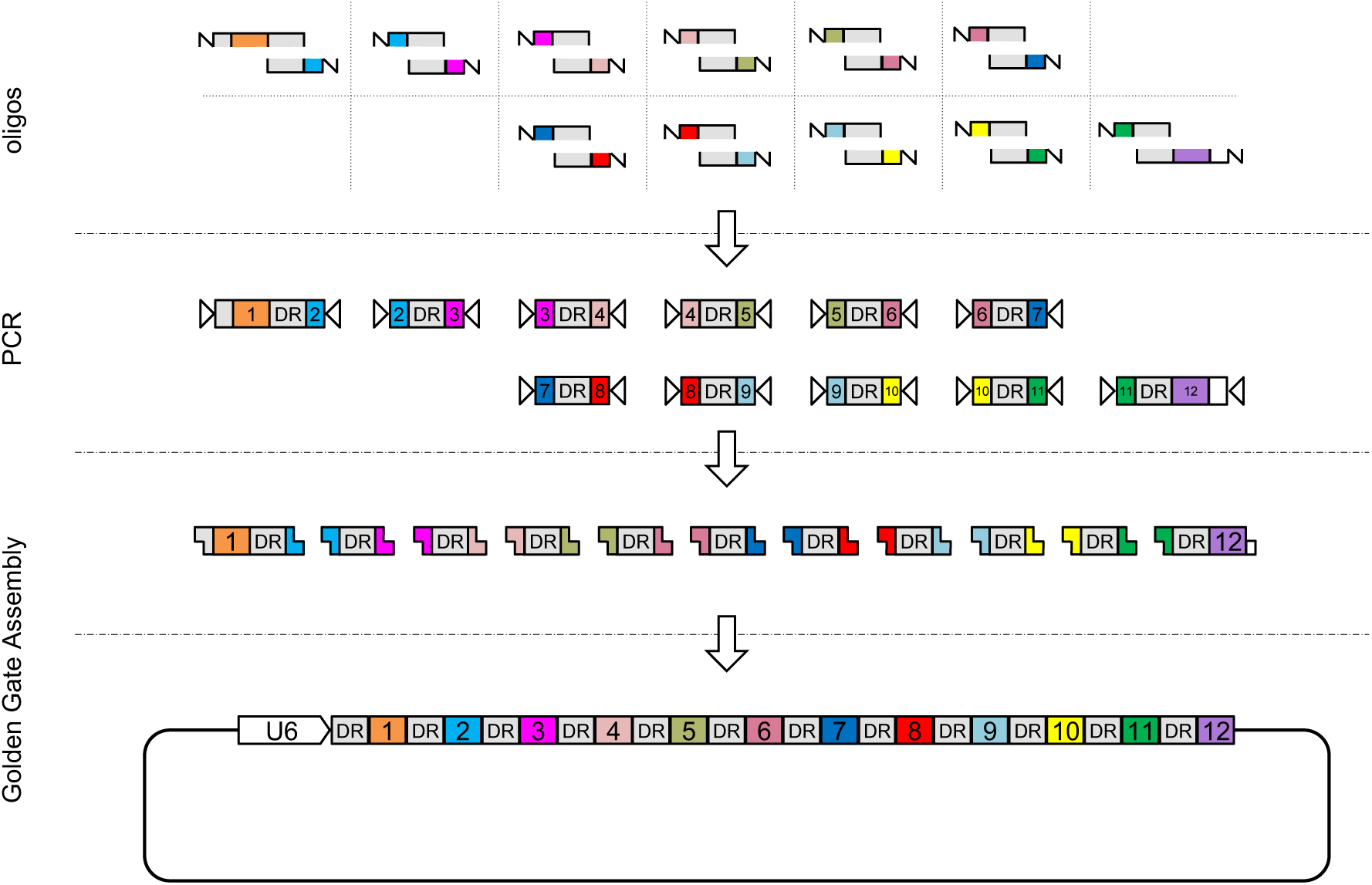

## Supplementary Methods

### Protocol 1: Assembly of CRISPR arrays for (d)(en)AsCas12a with conventional Anneal-based strategies

Reagents: BsaI-HF v2 (NEB, R3733), T4 DNA Ligase (NEB, M0202).

1) Set up the annealing reactions (total volume of 20µl for each reaction) in PCR tubes as follows:

2) Transfer PCR tubes to a thermocycler and run the following annealing program:

| component | volume | final concentration |
| --- | --- | --- |
| forward primer (10 µM) | 5 µl | 2.5 µM |
| reverse primer (10 µM) | 5 µl | 2.5 µM |
| T4 DNA Ligase Buffer (10×) | 2 µl | 1× |
| ddH<sub>2</sub>O | 8 µl |  |

3) Dilute annealed oligos to a final concentration of 0.5 µM (i.e., 5-fold dilution by adding 80 µl ddH_2_O to each reaction).
4) Three different versions of assembly were performed individually as indicated in Supplementary Figure S4: (-)BsaI, (+)BsaI, Gold Gate.
  4.1) Prepare the assembly reaction:

- 2) Transfer PCR tubes to a thermocycler and run the following annealing program:
| step | temperature | duration | cycles |
| --- | --- | --- | --- |
| 1 | 95°C | 4 min |  |
| 2 | 70°C | 10 min |  |
| 3 | 70°C (set: -0.1°C/cycle) | 2 sec |  |
| 4 | go to step 3 |  | 500 |
| end |  |  |  |

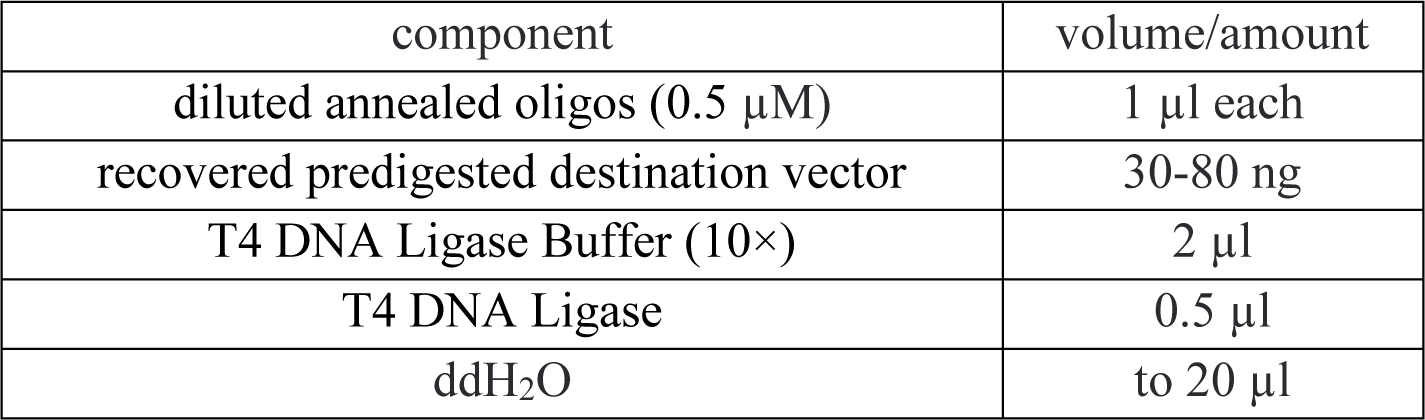 Incubate at 16°C overnight with a thermocycler.
  4.2) Prepare the assembly reaction:

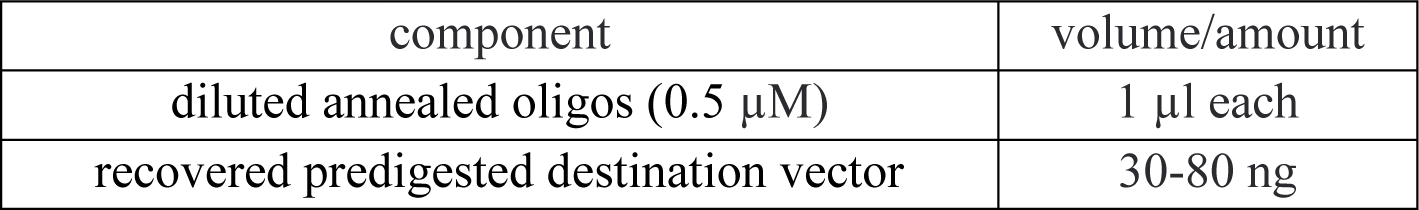
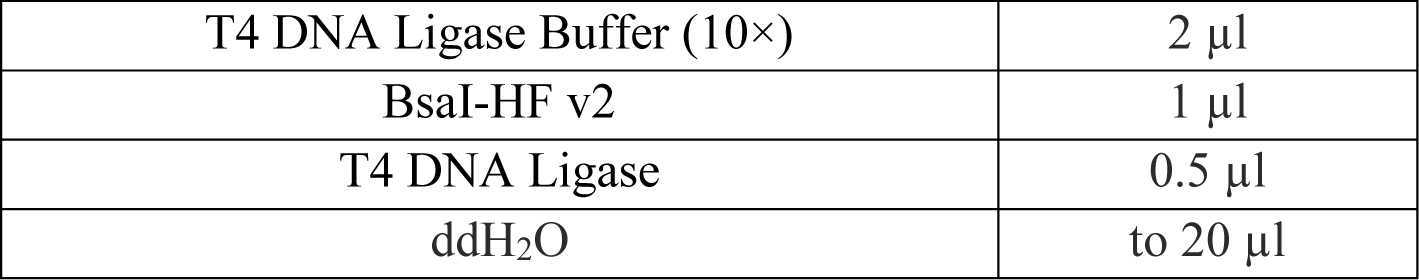 Incubate at 16°C overnight with a thermocycler, followed by 37°C for 15 min before transformation.
  4.3) Prepare the assembly reaction:

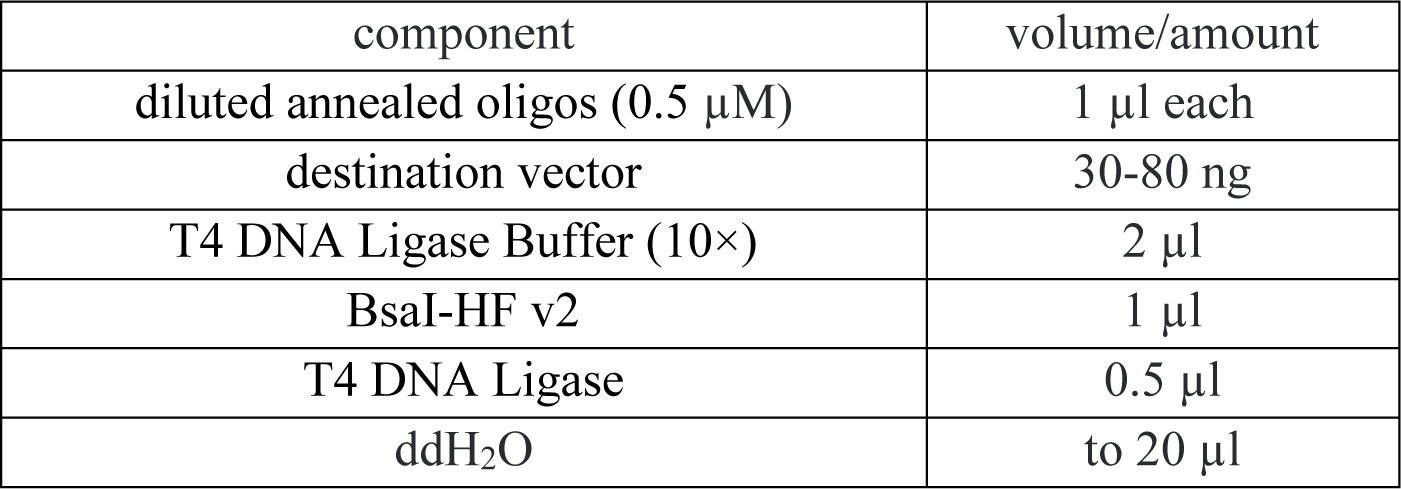 Run the following Golden Gate Assembly program with a thermocycler: (37°C 5 min → 16°C 5 min) × 30 cycles, followed by 60°C for 5 min. If reactions are done overnight, add a 4°C terminal hold to the program and repeat the step of 60°C for 5 min the next day before transformation.
5) Transform 2-10 µl of the assembly reaction into competent cells.

### Protocol 2: Assembly of CRISPR arrays for (d)(en)AsCas12a with PCR-based strategy

Reagents: BsaI-HF v2 (NEB, R3733), T4 DNA Ligase (NEB, M0202), Taq DNA Polymerase (2 ×premix), sodium acetate (3M, pH 5.2).

1) Set up the PCR reactions (total volume of 50µl for each reaction) as follows:

2) Transfer PCR tubes to a thermocycler and run a routine PCR program:

| component | volume | final concentration |
| --- | --- | --- |
| forward primer (10 µM) | 5 µl | 1 µM |
| reverse primer (10 µM) | 5 µl | 1 µM |
| Taq premix (2 ×) | 25 µl | 1 × |
| ddH<sub>2</sub>O | 15 µl |  |

3) Upon completion, transfer PCR products to 1.5 ml tubes. Add 1/10 volume of NaAc (3M, pH 5.2), and 2 volumes of pre-chilled 100% ethanol. Mix well. Store at -20°C for 20 min to overnight to precipitate the DNA.
4) Centrifuge at 10,000-20,000g for 15 min (at 4°C if convenient). Carefully pour off the ethanol (or remove by pipetting) without disturbing the pellet. Wash twice with 70-75% ethanol. Allow the DNA pellet to air-dry. Dissolve the DNA 20 µl TE buffer or ddH_2_O.
5) Quantify the concentration of recovered DNA samples. Adjust them to a uniform concentration (e.g., around 5 ng/µl).
6) Prepare the assembly reaction:

- 2) Transfer PCR tubes to a thermocycler and run a routine PCR program:
| step | temperature | duration | cycles |
| --- | --- | --- | --- |
| 1 | 95°C | 3 min |  |
| 2 | 95°C | 30 sec |  |
| 3 | 45-50°C | 30 sec |  |
| 4 | 72°C | 30 sec |  |
| 5 | go to step 2 |  | 34 |
| 6 | 72°C | 5 min |  |

7) Run the following Golden Gate Assembly program with a thermocycler: (37°C 5 min → 16°C 5 min) × 30-60 cycles, followed by 60°C for 5 min. If reactions are done overnight, add a 4°C terminal hold to the program and repeat the step of 60°C for 5 min the next day before transformation.
8) Transform 2-10 µl of the assembly reaction into competent cells.

| component | volume/amount |
| --- | --- |
| recovered PCR segments | ~5 ng each |
| destination vector | 30-80 ng |
| T4 DNA Ligase Buffer (10 ×) | 2 µl |
| BsaI-HF v2 | 1-2 µl |
| T4 DNA Ligase | 0.5 µl |
| ddH<sub>2</sub>O | to 20 µl |

### Protocol 3: A simplified version of Protocol 2

Reagents: BsaI-HF v2 (NEB, R3733), T4 DNA Ligase (NEB, M0202), Taq DNA Polymerase (2 ×premix).

Note: The optimal primer concentration for PCR reaction and the volume pipetted for subsequent assembly were determined based on the designated PCR reagent used in this study. Although these reaction parameters may need to be optimized for different PCR reagents, they can be used as benchmarks. When necessary, adjust them up or down to obtain satisfactory results.

1) Set up the PCR reactions (total volume of 20µl for each reaction) as follows:

2) Transfer PCR tubes to a thermocycler and run a routine PCR program:

| component | volume | final concentration |
| --- | --- | --- |
| forward primer (10 µM) | 5 µl | 2.5 µM |
| reverse primer (10 µM) | 5 µl | 2.5 µM |
| Taq premix (2×) | 10 µl | 1 × |

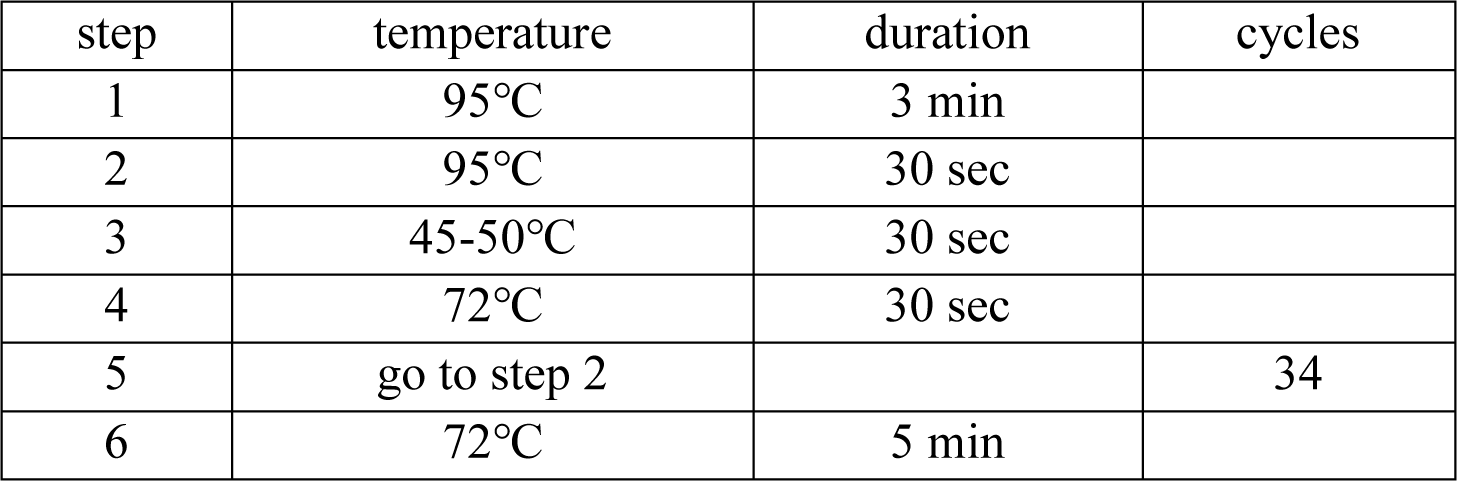
3) Upon completion, dilute PCR products 10-fold by adding 180 µl ddH_2_O to each reaction.
4) Prepare the assembly reaction:

5) Run the following Golden Gate Assembly program with a thermocycler: (37°C 5 min → 16°C 5 min) × 30-60 cycles, followed by 60°C for 5 min. If reactions are done overnight, add a 4°C terminal hold to the program and repeat the step of 60°C for 5 min the next day before transformation.
6) Transform 2-10 µl of the assembly reaction into competent cells.

- 3) Upon completion, dilute PCR products 10-fold by adding 180 µl ddH<sub>2</sub>O to each reaction. - 4) Prepare the assembly reaction:
| component | volume/amount |
| --- | --- |
| diluted PCR products | 1 µl each |
| destination vector | 30-80 ng |
| T4 DNA Ligase Buffer (10×) | 2 µl |
| BsaI-HF v2 | 1-2 µl |
| T4 DNA Ligase | 0.5 µl |
| ddH<sub>2</sub>O | to 20 µl |

### Protocol 4: Assembly of CRISPR arrays for RfxCas13d with PCR-based strategy

Reagents: BsaI-HF v2 (NEB, R3733), T4 DNA Ligase (NEB, M0202), PrimeSTAR HS DNA Polymerase (Takara, R010A).

1) Set up the PCR reactions (total volume of 50µl for each reaction) as follows:

2) Transfer PCR tubes to a thermocycler and run a routine PCR program:

| component | volume/amount | final concentration |
| --- | --- | --- |
| 5 × Buffer (Mg<sup>2+</sup> Plus) | 10 µl | 1 × |
| dNTP mixture (2.5 mM each) | 4 µl | 200 µM each |
| DNA polymerase | 0.5 µl |  |
| forward primer (10 µM) | 5 µl | 1 µM |
| reverse primer (10 µM) | 5 µl | 1 µM |
| template plasmid | 0.2-1 ng |  |
| ddH<sub>2</sub>O | to 50 µl |  |

3) Upon completion, purify PCR segments with available DNA recovery kit. Quantify the concentration. Adjust them to a uniform concentration (e.g., around 5 ng/µl).
4) Prepare the assembly reaction:

- 2) Transfer PCR tubes to a thermocycler and run a routine PCR program:
| step | Temperature | duration | cycles |
| --- | --- | --- | --- |
| 1 | 98°C | 30 sec |  |
| 2 | 98°C | 10 sec |  |
| 3 | 50°C | 15 sec |  |
| 4 | 72°C | 30 sec |  |
| 5 | go to step 2 |  | 34 |
| 6 | 72°C | 5 min |  |

5) Run the following Golden Gate Assembly program with a thermocycler: (37°C 5 min → 16°C 5 min) × 30-60 cycles, followed by 60°C for 5 min. If reactions are done overnight, add a 4°C terminal hold to the program and repeat the step of 60°C for 5 min the next day before transformation.
6) Transform 2-10 µl of the assembly reaction into competent cells.

| component | volume/amount |
| --- | --- |
| recovered PCR segments | ~5 ng each |
| destination vector | 30-80 ng |
| T4 DNA Ligase Buffer (10×) | 2 µl |
| BsaI-HF v2 | 1-2 µl |
| T4 DNA Ligase | 0.5 µl |
| ddH<sub>2</sub>O | to 20 µl |

## Supplementary Tables

**Table S1.** Primers used for qPCR.

**Table S2.** Gene-specific primers used for reverse transcription (also used as the reverse primer of subsequent qPCR) and counterpart qPCR primers.

**Table S3.** Spacer sequences for transcriptional activation mediated by dAsCas12a-VPR or denAsCas12a-VPR.

**Table S4.** Spacer sequences for RfxCas13d-mediated RNA cleavage.

**Table S5.** Spacer sequence for gene editing mediated by enAsCas12a.

**Table S6.** Primers used for the amplification and sequencing of the target region flanking indels.

**Table S7.** Primers used to assemble the CRISPR array of 6 crRNAs for dAsCas12a-VPR or denAsCas12a-VPR with conventional Anneal-based strategies.

**Table S8.** Primers used to assemble the CRISPR array of 7 crRNAs for dAsCas12a-VPR or denAsCas12a-VPR with conventional Anneal-based strategies.

**Table S9.** Primers used to assemble the CRISPR array of 6 crRNAs for dAsCas12a-VPR or denAsCas12a-VPR with PCR-based strategy.

**Table S10.** Primers used to assemble the CRISPR array of 7 crRNAs for dAsCas12a-VPR or denAsCas12a-VPR with PCR-based strategy.

**Table S11.** Primers used to assemble the CRISPR array of 9 crRNAs for dAsCas12a-VPR or denAsCas12a-VPR with PCR-based strategy.

**Table S12.** Primers used to assemble the CRISPR array of 12 crRNAs for dAsCas12a-VPR or denAsCas12a-VPR with PCR-based strategy.

**Table S13.** Primers used to assemble 12 crRNAs for dAsCas12a-VPR or denAsCas12a-VPR based on mutant DR.

**Table S14.** Primers used to assemble the CRISPR array of 9 crRNAs for RfxCas13d with PCR-based strategy.

**Table S15.** Primers used to assemble the CRISPR array of 12 crRNAs for RfxCas13d with PCR-based strategy.

**Table S16.** Primers used to assemble the CRISPR array of 15 crRNAs for RfxCas13d with PCR-based strategy.

**Table S17.** Primers used to assemble 12 crRNAs for dAsCas12a-VPR or denAsCas12a-VPR with two rounds of assembly.

**Table S18.** Primers used to assemble 20 crRNAs for dAsCas12a-VPR or denAsCas12a-VPR with two rounds of assembly.

**Table S19.** Primers used to assemble U6-4-U6-4-U6-4 for dAsCas12a-VPR or denAsCas12a-VPR with two rounds of assembly.

**Table S20.** Primers used to assemble U6-4-U6-4-U6-4-U6-3 for RfxCas13d with two rounds of assembly.

**Table S21.** Sequences of constructs used in this study.

## Supplementary Figures

**Figure S1.**
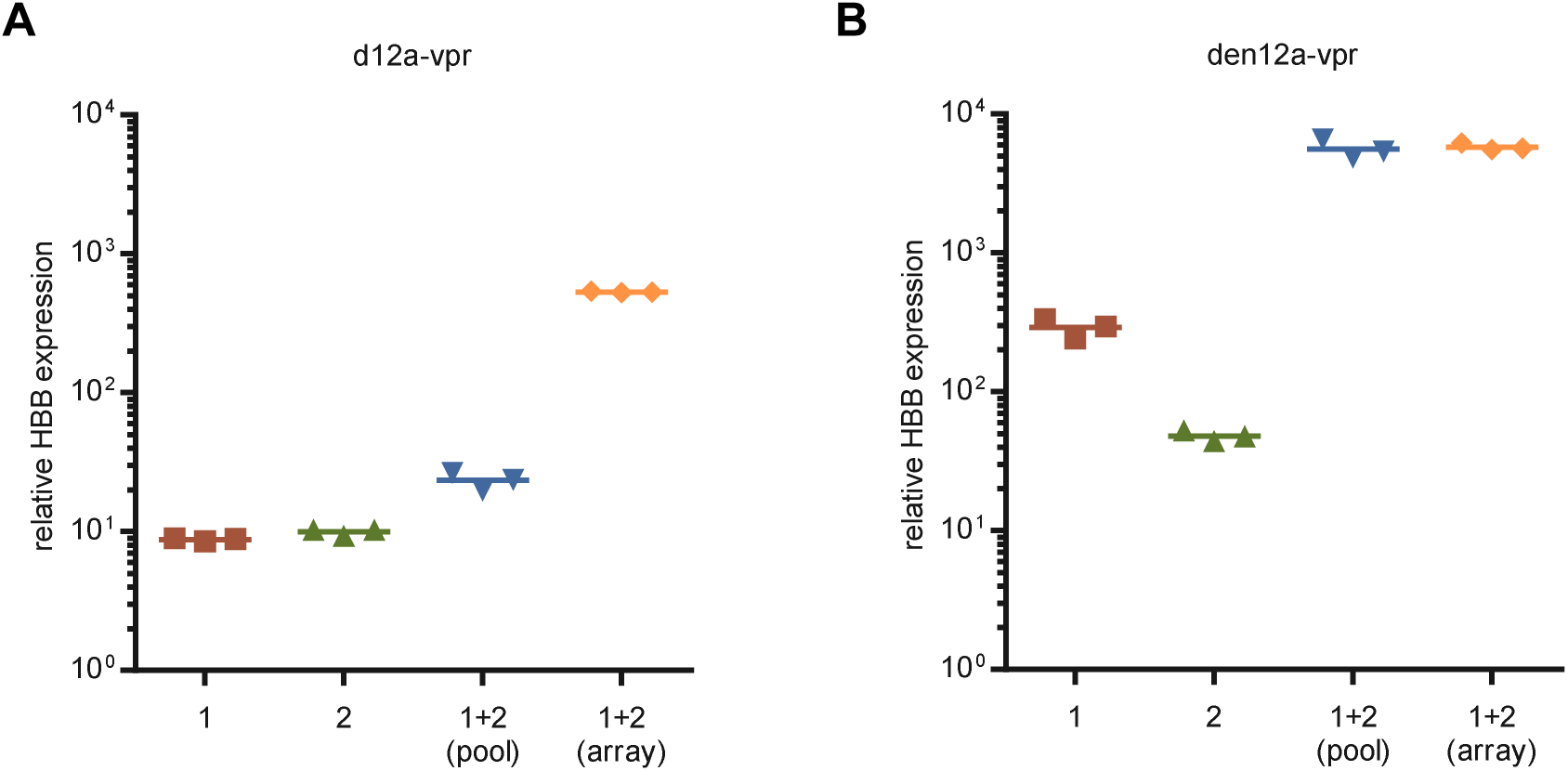
Synergistic transcriptional activation of *HBB* with d12a-VPR or den12a-VPR. **(A**, **B)** Quantification of relative *HBB* expression over non-targeting control in HEK293T cells 48h after transfection with plasmids encoding d12a-VPR **(A)** or den12a-VPR **(B)** and *HBB* promoter-targeting crRNA-1, crRNA-2, a pool of both, or an array containing both. Values shown as mean, n = 3 independent experiments.

**Figure S2.**
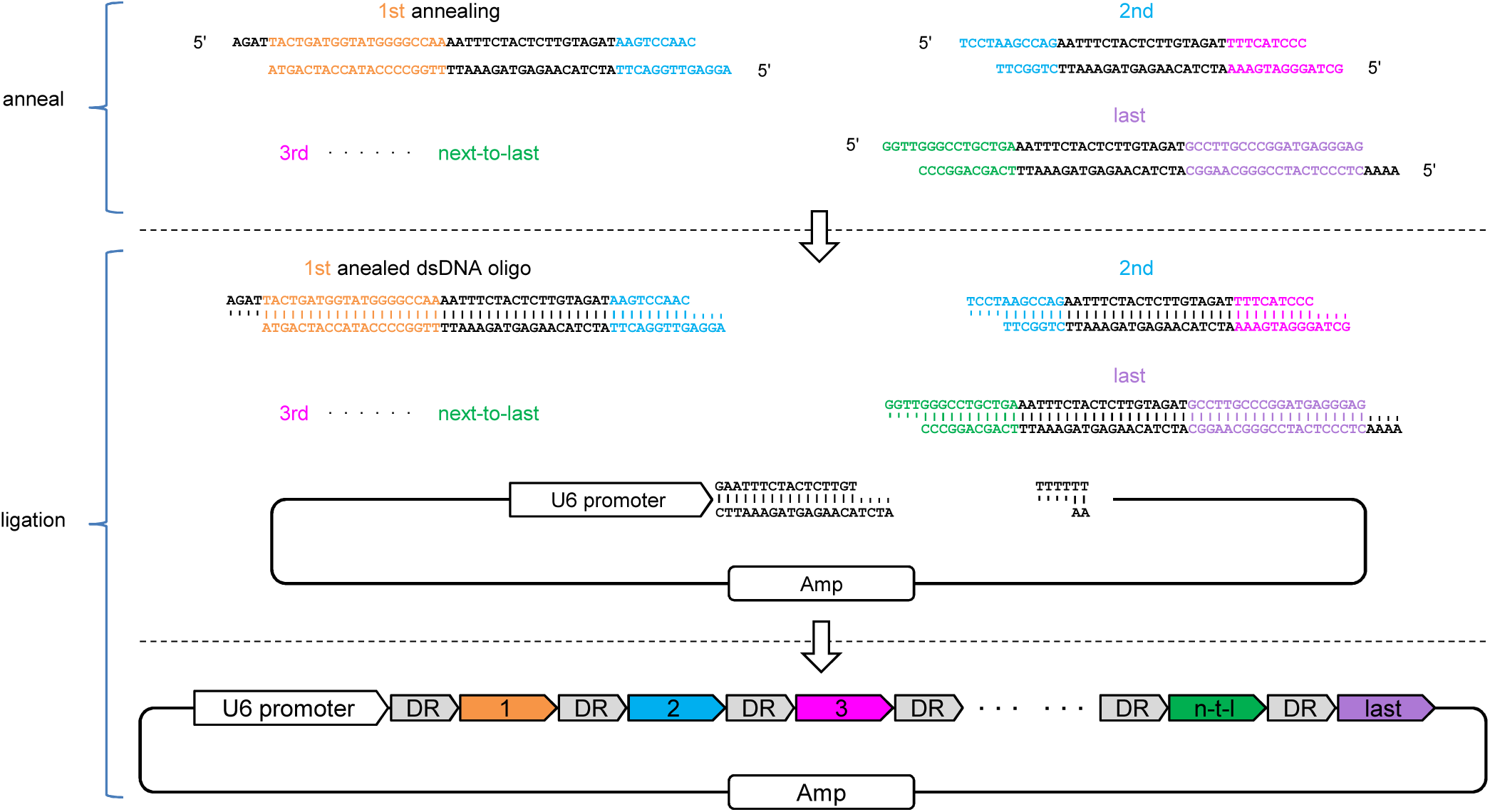
Schematic illustration and workflow of conventional Anneal-based CRISPR array assembly strategy. First, anneal single-stranded oligo pairs to form double-stranded DNA segments with desired sticky ends. Then, set up and run a ligation reaction with diluted annealed oligos and destination cloning vector (with/without predigestion and purification). Detailed protocols are provided in Supplementary Methods.

**Figure S3.**
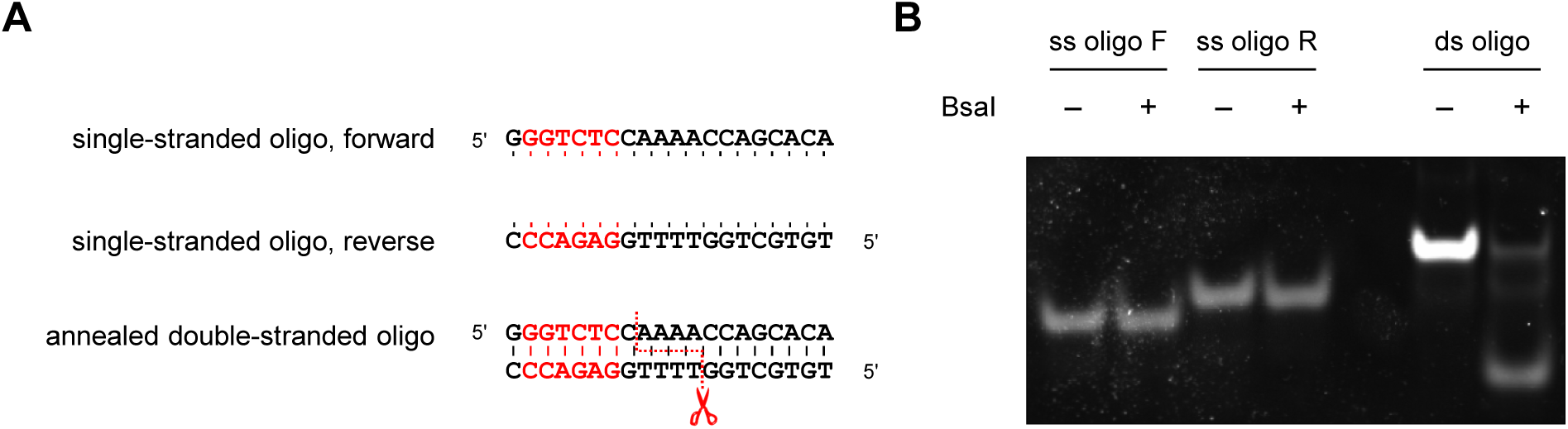
Restriction endonuclease BsaI specifically cleaves double-stranded DNA substrates. **(A)** Reverse complementary single-stranded oligos and resulting annealed double-stranded product. Recognition site of BsaI is colored red. Cleavage site is indicated by red dashed line. **(B)** Representative PAGE image of oligos depicted in (**A**) after incubation with/without BsaI at 37℃ for 1 hour.

**Figure S4.**
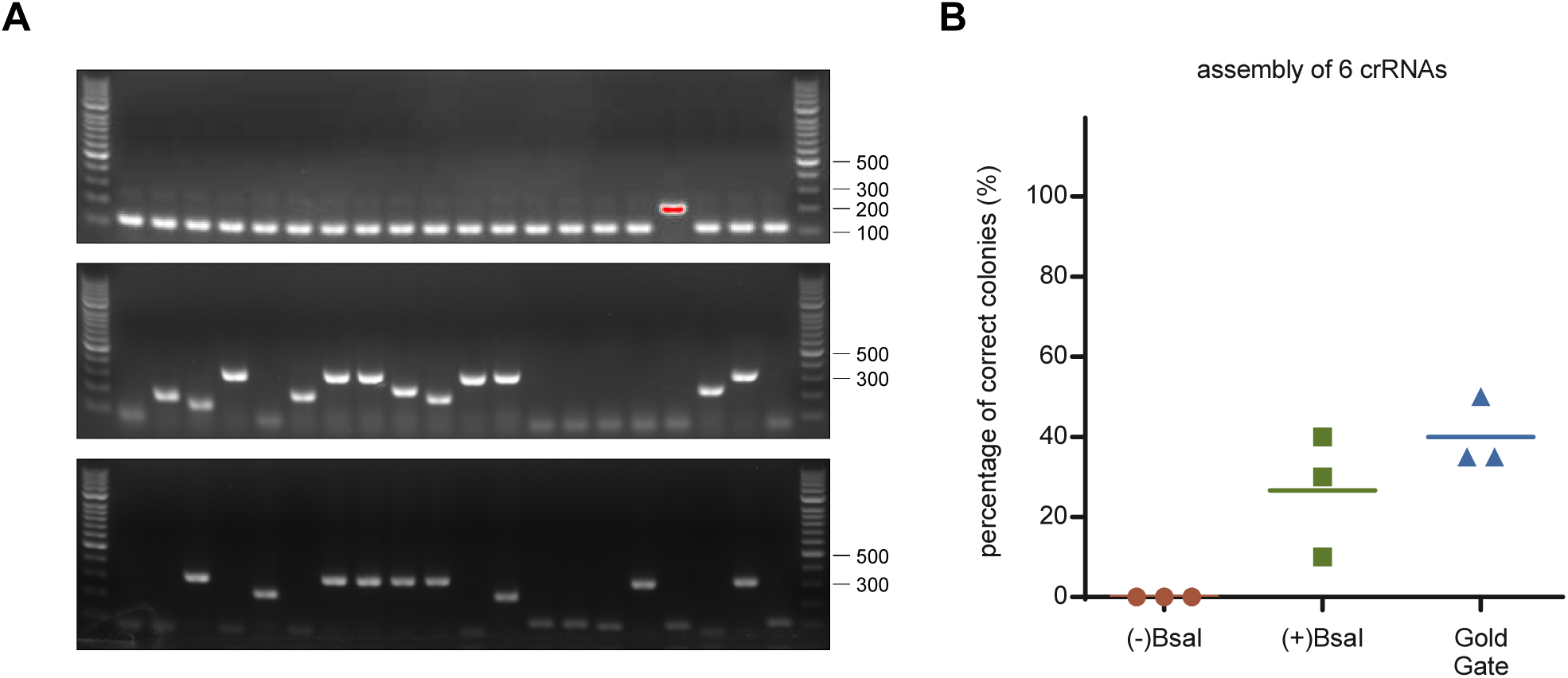
Comparison of assembly accuracy across different versions of conventional Anneal-based strategy. **(A)** Representative images of colony PCR to evaluate the accuracy of assembling 6 crRNAs using three versions of Anneal-based strategy: (-)BsaI (upper), (+)BsaI (middle), Gold Gate (lower), and the corresponding accuracies are shown in **(B)**. (-)BsaI: pre-digested cloning vector is recovered and added to a ligation reaction with annealed oligos, incubated overnight at room temperature, then directly transformed into competent cells; (+)BsaI: same as (-)BsaI, except that the ligation mixture is re-digested with BsaI for 15 minutes before transformation; Gold Gate: circular cloning vector and annealed oligos are added to a Gold Gate reaction, and then run a standard Gold Gate assembly program (37℃ 5 min → 16℃ 5 min, 30 cycles, followed by 60℃ 5 min) with a thermocycler. Detailed protocols are provided in Supplementary Methods. Values shown as mean, n = 3 independent experiments.

**Figure S5.**
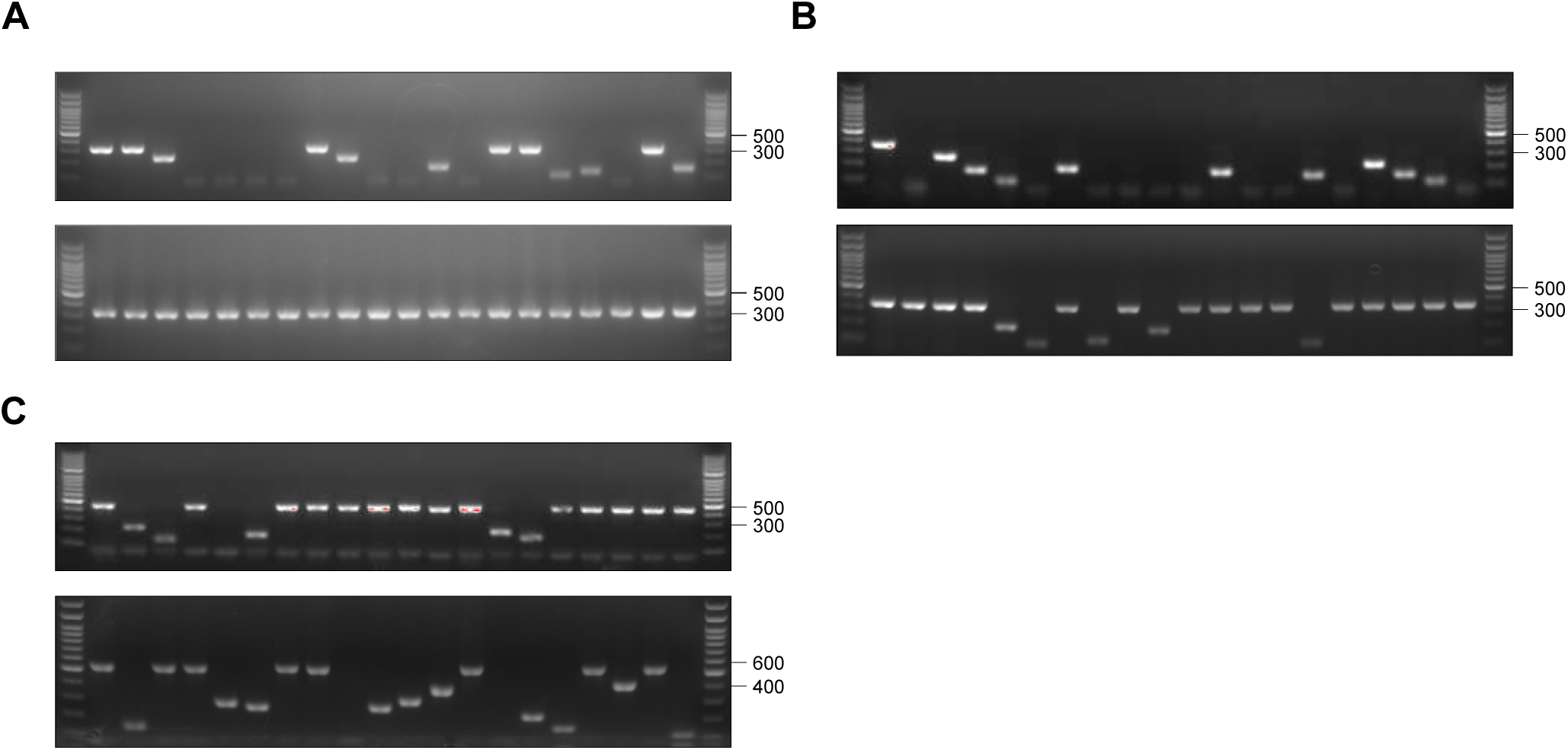
High-accuracy assembly of CRISPR array with PCR-based strategy. (**A**, **B**) Representative images of colony PCR to evaluate the accuracy of assembling 6 (**A**) or 7 (**B**) crRNAs using conventional Anneal-based (upper) or PCR-based strategy (lower). (**C**) Representative images of colony PCR to evaluate the accuracy of assembling 9 (upper) or 12 (lower) crRNAs using PCR-based strategy.

**Figure S6.**
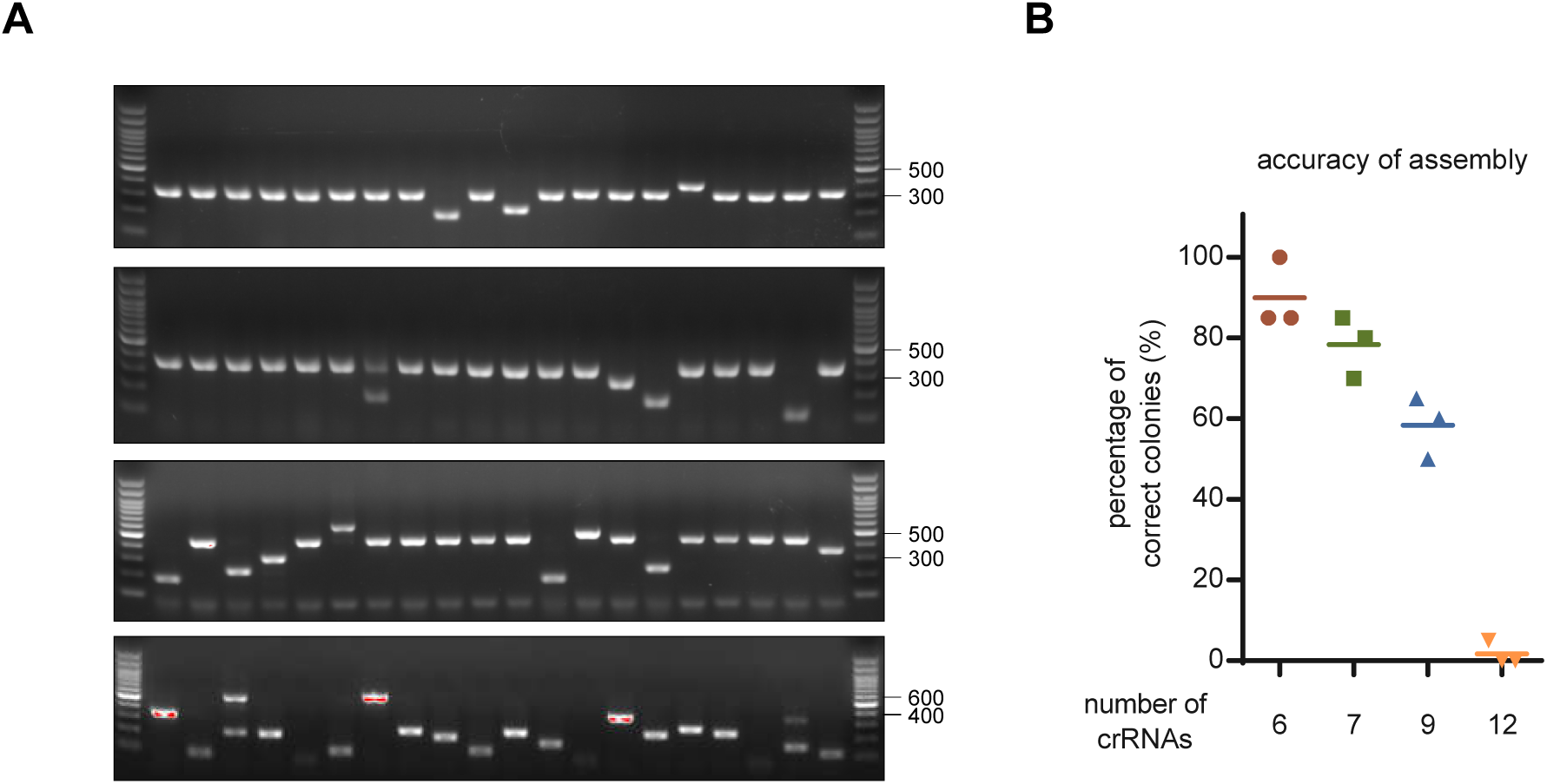
Accuracies of assembling 6 to 12 crRNAs using simplified recovery-free version of PCR-based strategy. (**A**) Representative images of colony PCR when assembling 6 (upper), 7 (middle-upper), 9 (middle-lower), and 12 (lower) crRNAs, and the corresponding accuracies are shown in **(B)**. Values shown as mean with n = 3.

**Figure S7.**
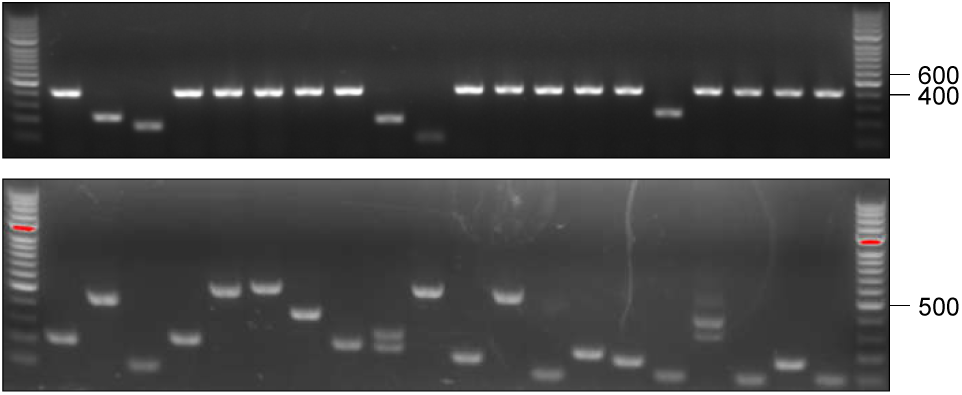
Representative images of colony PCR when assembling 9 (upper) and 12 (lower) crRNAs using mutant DR with higher GC content.

**Figure S8.**
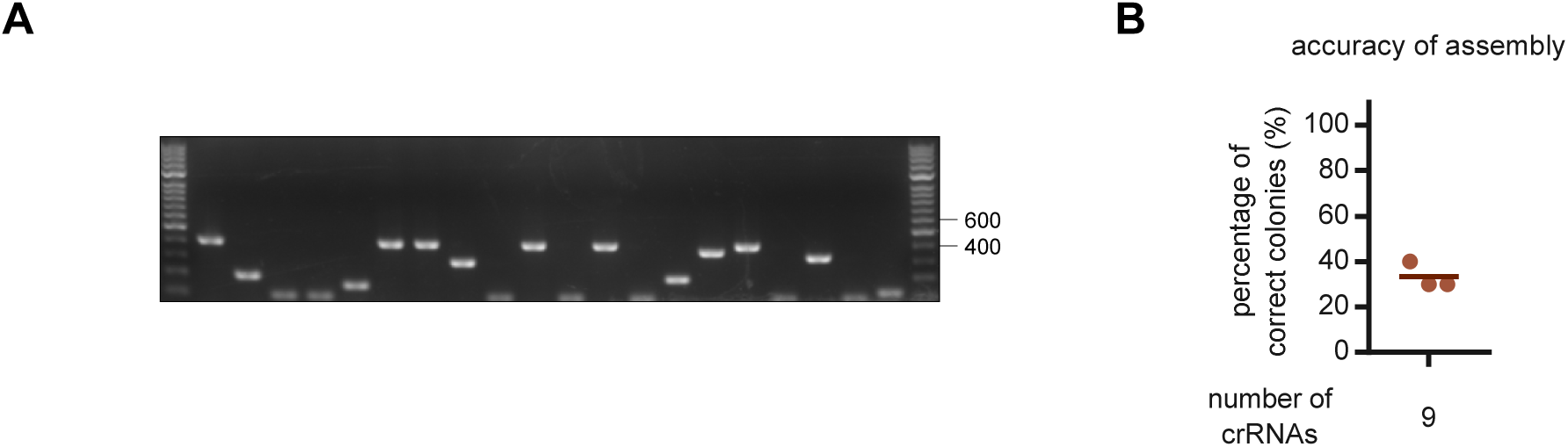
Accuracies of assembling 9 crRNAs using simplified recovery-free version of PCR-based strategy and mutant DR with higher GC content. (**A**) Representative image of colony PCR, and the corresponding accuracies are shown in **(B)**. Values shown as mean with n = 3.

**Figure S9.**
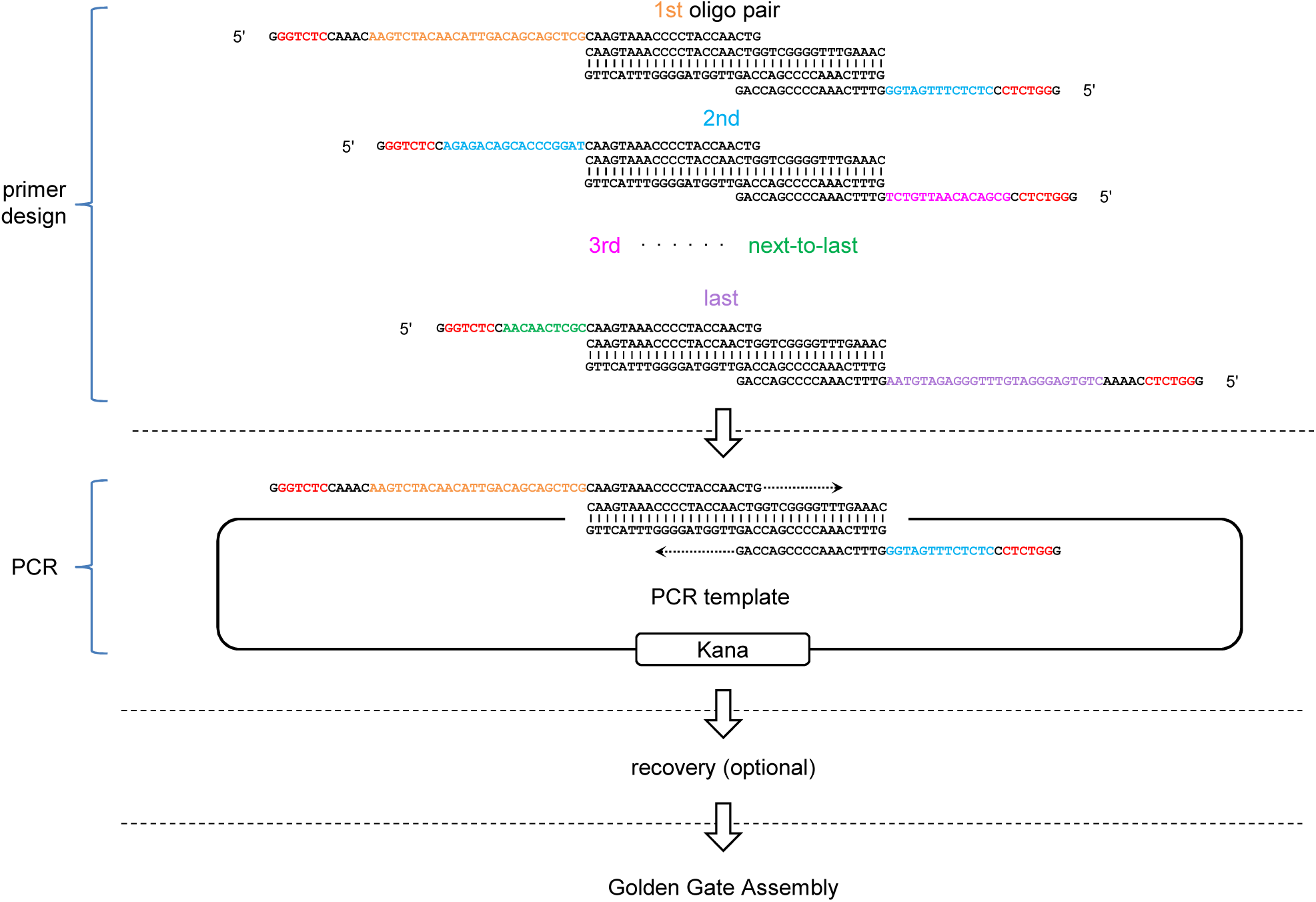
Schematic illustration and workflow of assembling crRNAs for RfxCas13d. Similar to Figure 1 except for the following minor modifications: (1) Primer pairs are designed to be partially (∼20 bases 3’ section) complementary to the 36bp DR of RfxCas13d, instead of to each other; (2) To further shorten the initial oligos for cost-saving, a kanamycin-resistant plasmid containing the 36nt DR of RfxCas13d, which could be easily home-made, is introduced as a template for PCR reactions; (3) The relatively long length of the PCR products (70∼100nt) renders them compatible with common commercial DNA recovery kits, which are therefore used to obtain purer dsDNA segments.

**Figure S10.**
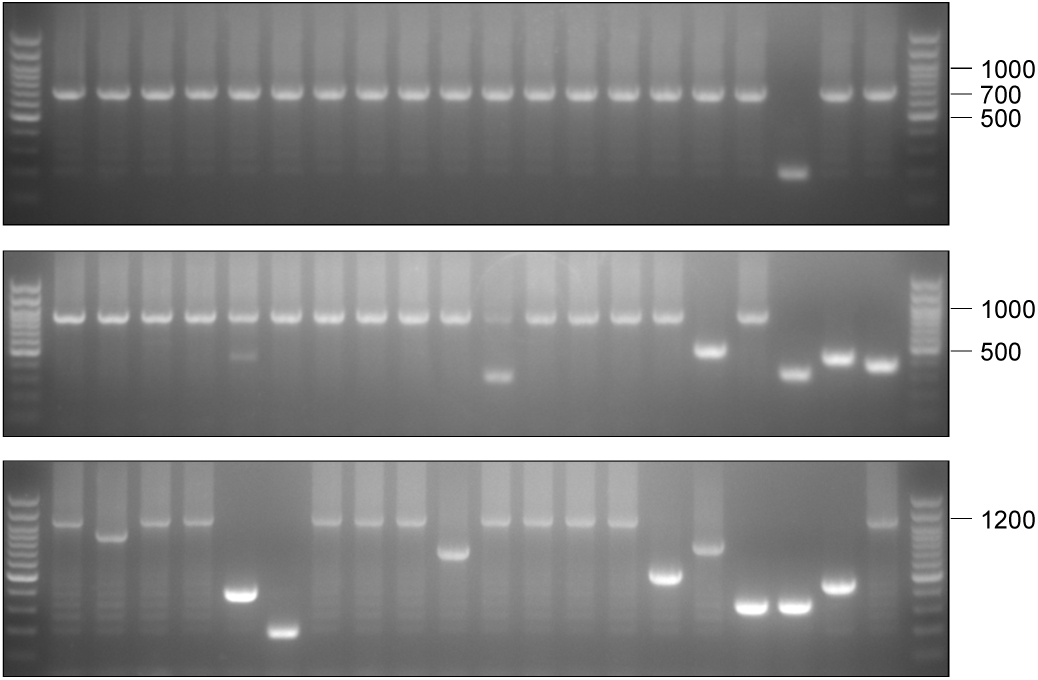
High-accuracy assembly of crRNAs for a Cas13d nuclease. Representative images of colony PCR to evaluate the accuracy of assembling 6 (upper), 9 (middle), or 12 (lower) crRNAs for RfxCas13d using PCR-based strategy.

**Figure S11.**
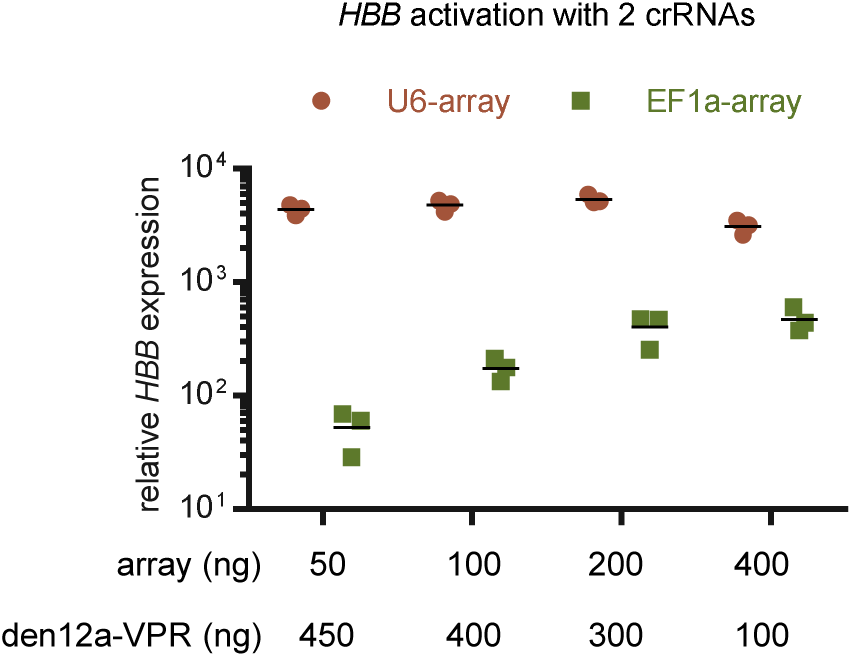
Quantification of relative *HBB* expression over non-targeting control in HEK293T cells 48h after transfection with the indicated amount of denAsCas12a-VPR and an array of 2 crRNAs targeting *HBB* promoter. Values shown as mean with n = 3.

**Figure S12.**
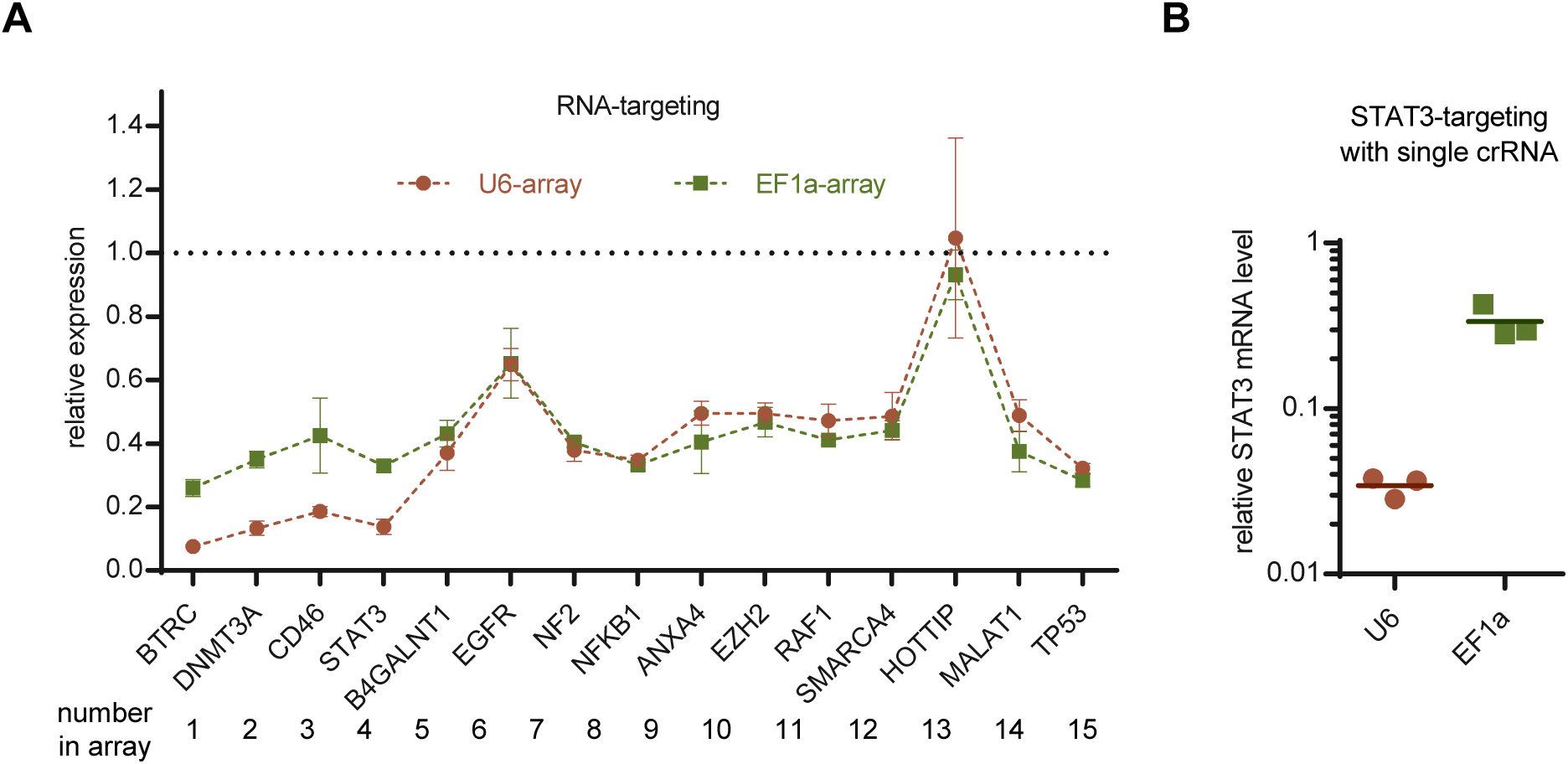
Distinct targeting patterns of Pol II/III promoter-driven CRISPR arrays with RfxCas13d system. (**A**) Quantification of relative mRNA expression of the indicated genes compared to non-targeting control in HEK293T cells 48h after transfection with RfxCas13d and an array of 15 crRNAs driven by either U6 (U6-array) or EF1a (EF1a-array). Values shown as mean ± sd with n = 3. (**B**) Quantification of relative STAT3 mRNA expression compared to non-targeting control in HEK293T cells 48h after transfection with RfxCas13d and a STAT3-targeting crRNA driven by either U6 or EF1a. Values shown as mean with n = 3.

**Figure S13.**
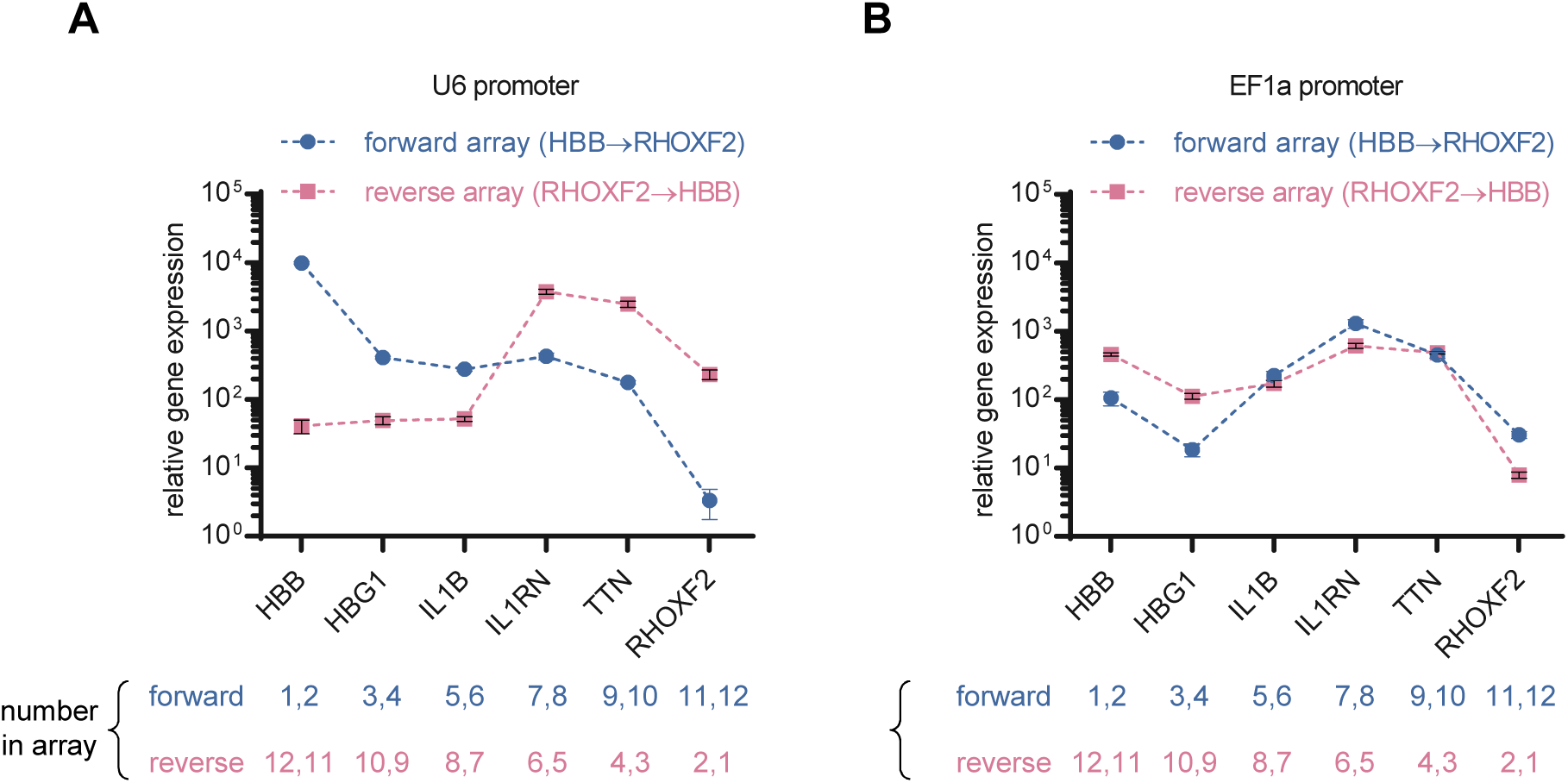
Transcription intensities across crRNAs located at different sites are milder but more evenly distributed in CRISPR arrays driven by Pol II promoters. (**A**) Quantification of relative mRNA expression over non-targeting control in HEK293T cells 48h after transfection with den12a-VPR and U6-driven arrays whose internal crRNAs are assembled in reverse order. (**B**) Quantification of relative mRNA expression over non-targeting control in HEK293T cells 48h after transfection with den12a-VPR and EF1a-driven arrays whose internal crRNAs are assembled in reverse order. Values shown as mean ± sd with n = 3.

**Figure S14.**
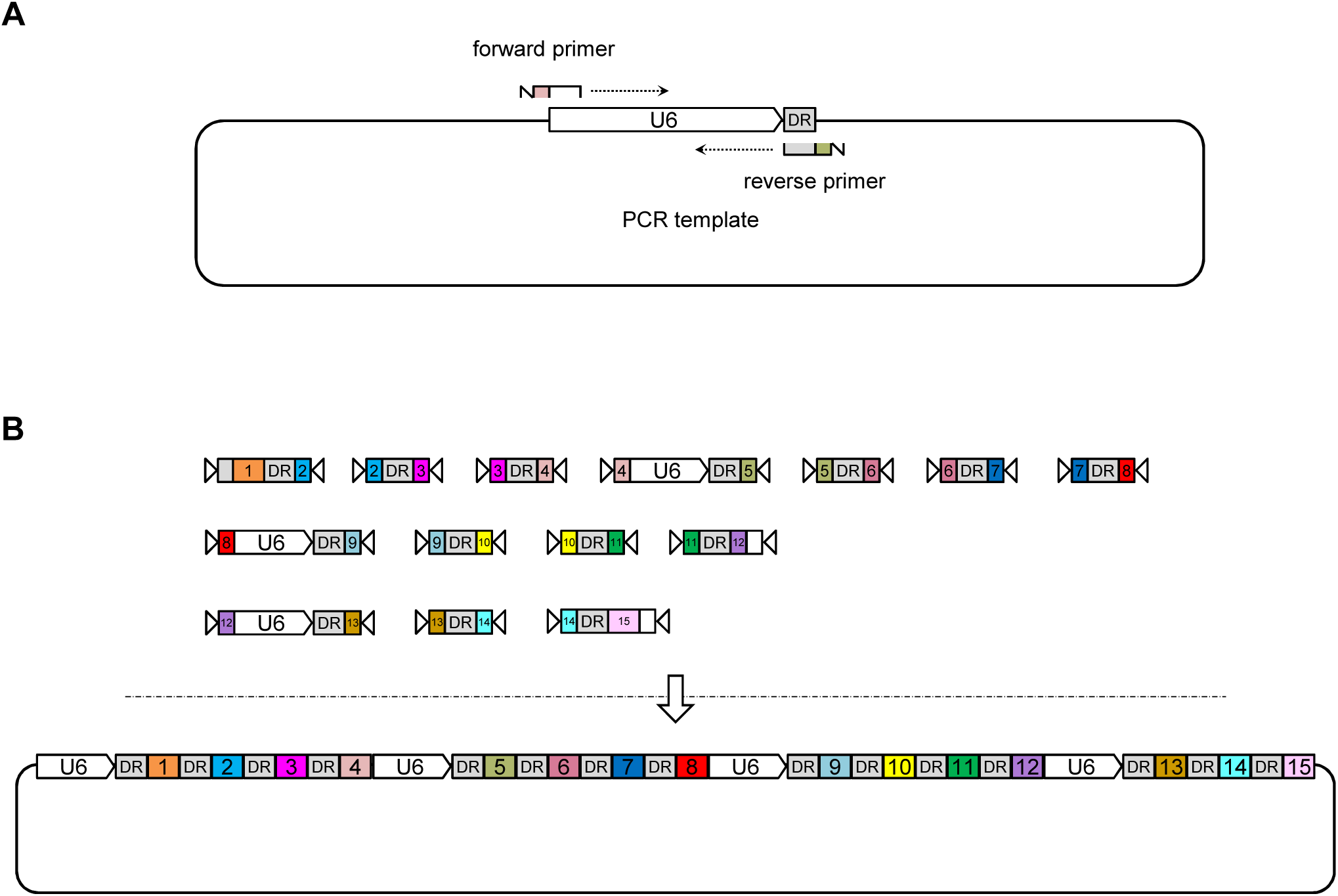
Assumption for the strategy to assemble a hierarchical CRISPR array. (**A**) Schematic illustration of the amplification of U6-containing fragments using a template containing a U6-DR cassette. (**B**) Schematic illustration of the proto assumption to assemble hierarchical CRISPR array.

**Figure S15.**
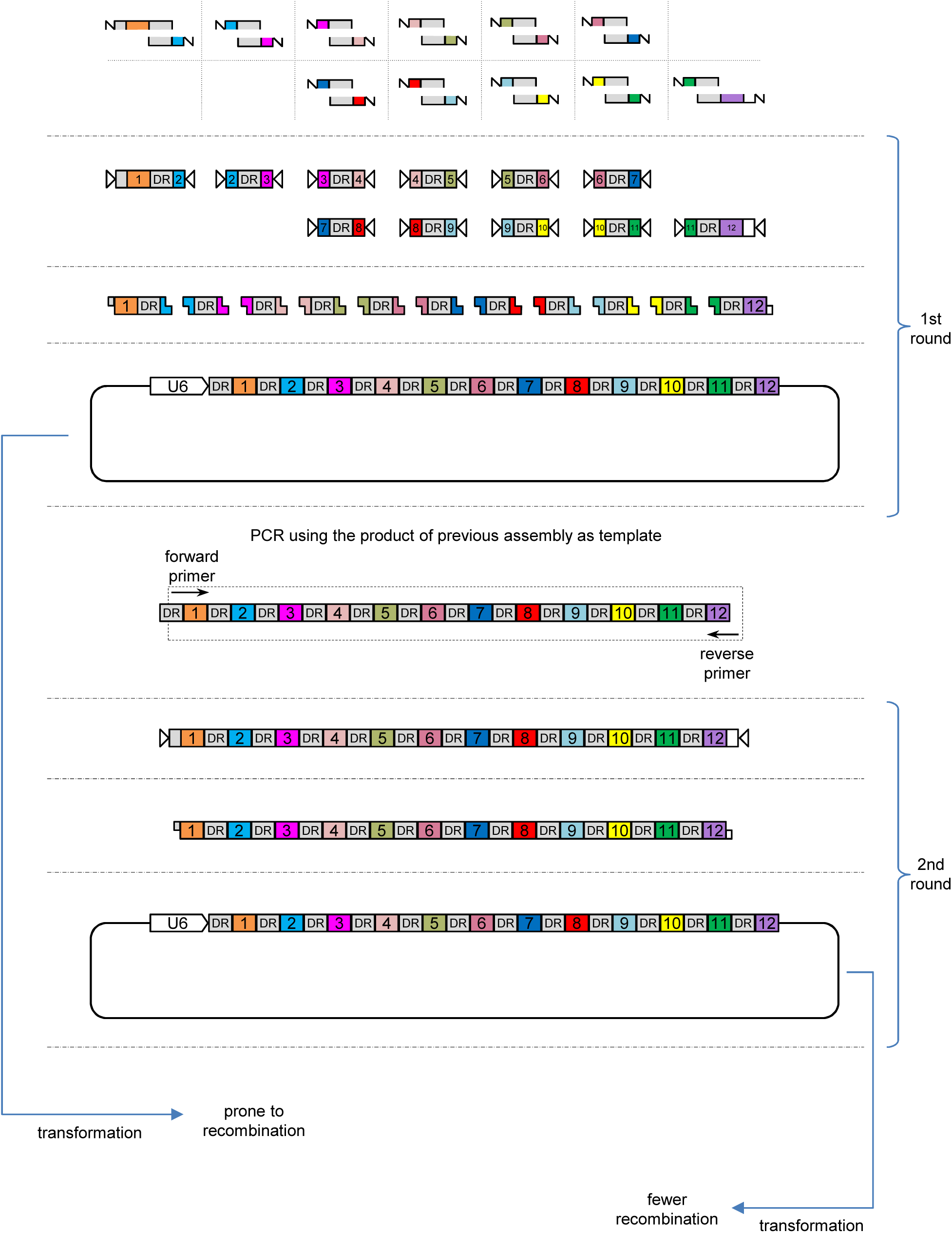
Schematic illustration and workflow of assembling CRISPR array using 2 rounds of assembly.

**Figure S16.**
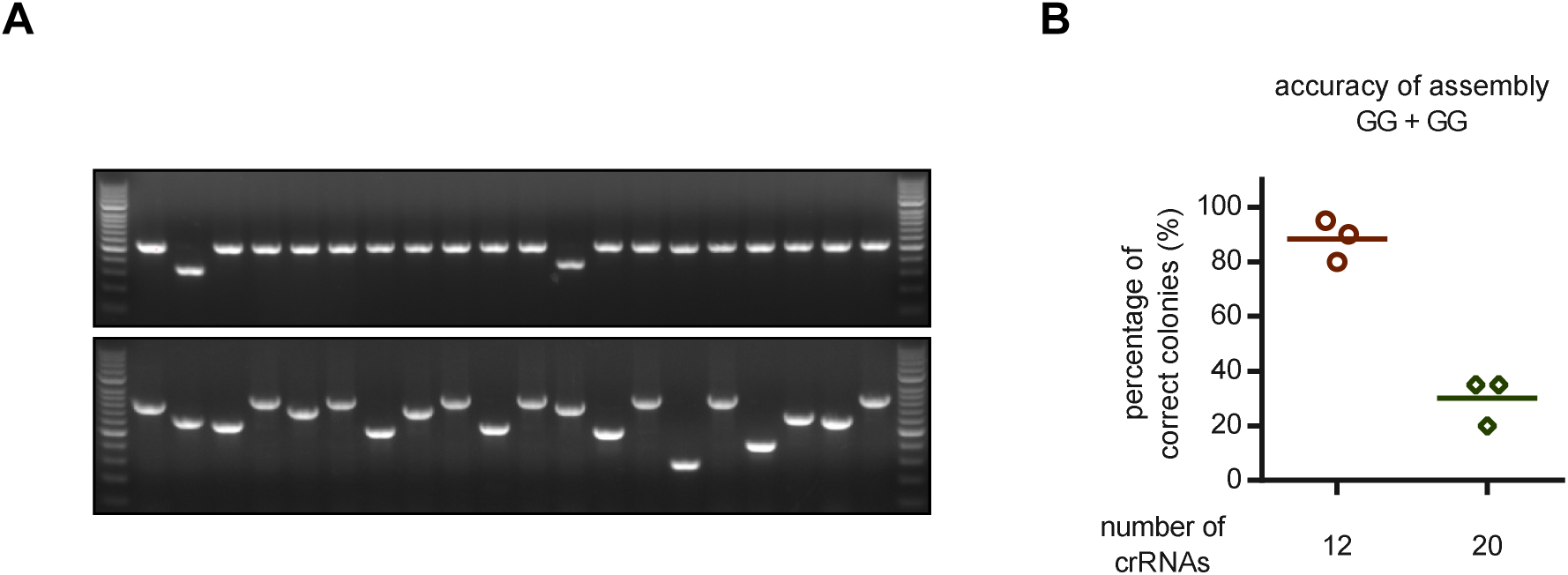
Accuracies of assembling 12 and 20 crRNAs using 2 rounds of assembly. (**A**) Representative images of colony PCR when assembling 12 (upper) and 20 (lower) crRNAs, and the corresponding accuracies are shown in (**B**). Note that since we failed to amplify the overall array from the assembly mixture of 20 crRNAs, the second round of PCR was separated into two individual reactions: crRNA 1-10, crRNA 11-20. Values shown as mean with n = 3.

**Figure S17.**
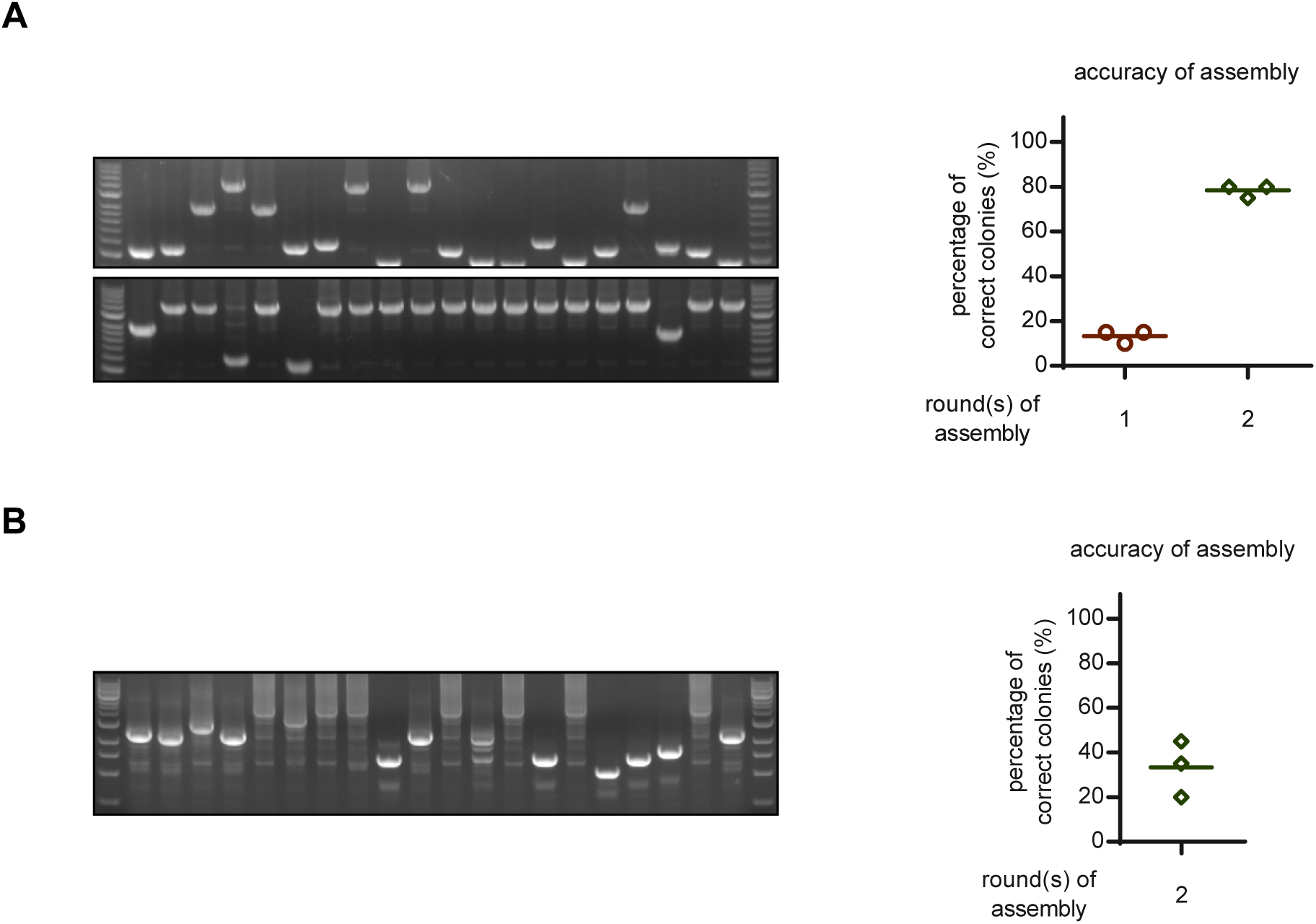
Assembly of complex, hierarchical CRISPR arrays. (**A**) Representative images of colony PCR when assembling an array consisting of 3 U6 promoters and 12 crRNAs, by either a single (upper) or two (lower) rounds of assembly, and the corresponding accuracies are shown in (right). (**B**) Representative image of colony PCR when assembling an array consisting of 4 U6 promoters and 15 crRNAs by two rounds of assembly, and the corresponding accuracies are shown in (right). Values shown as mean with n = 3.

**Figure S18.**
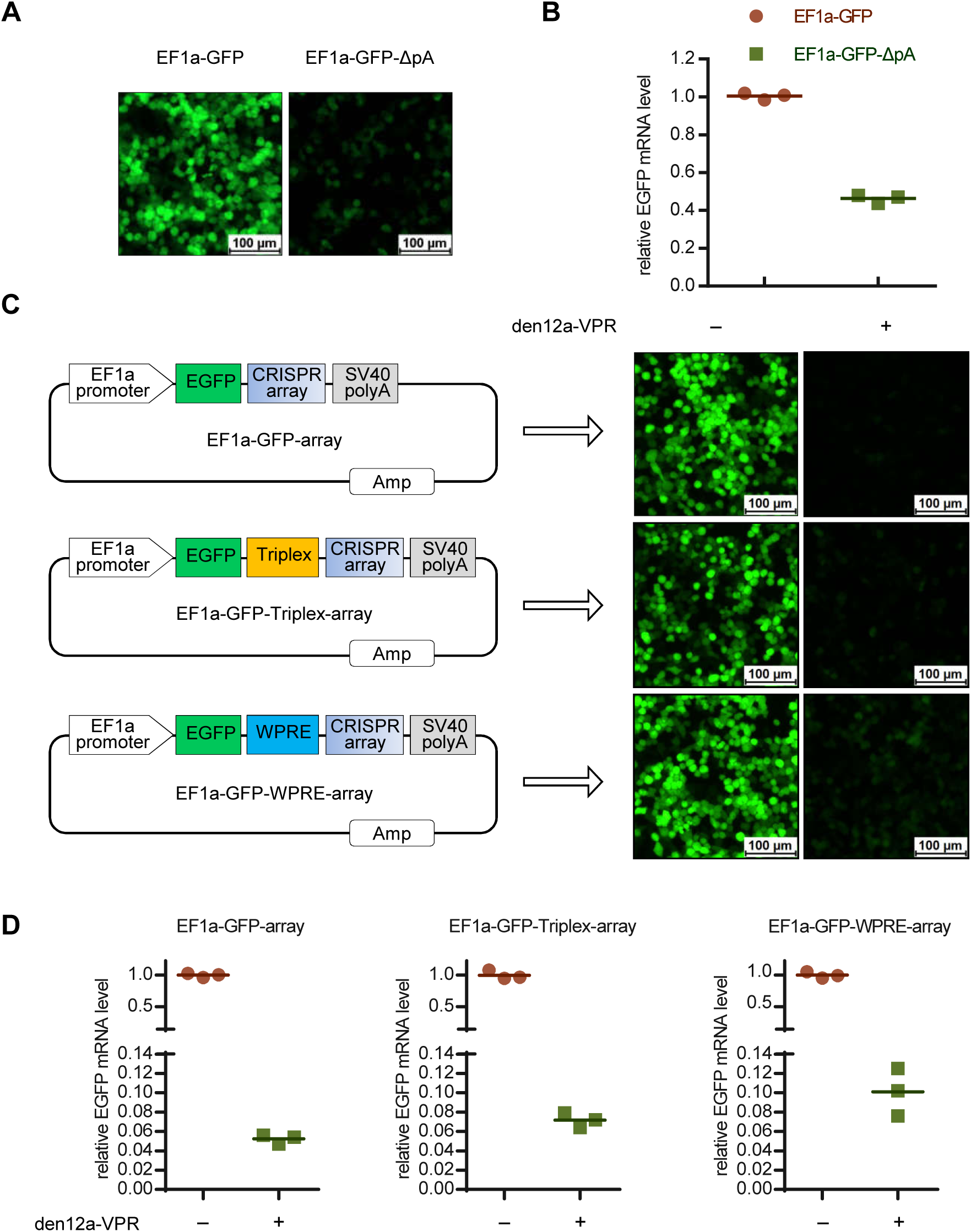
Processing of CRISPR array leads to inadequate expression of upstream EGFP encoded on the same transcript. **(A**, **B)** Representative fluorescence images **(A)** and mRNA expression **(B)** in HEK293T cells 48h after transfection with EF1a-driven EGFP constructs with/without a downstream SV40-poly(A) signal. Scale bar, 100 μm. **(C)** Schematics of EGFP constructs (left) and corresponding representative fluorescence images (right) 48h after transfection together with/without den12a-VPR. Scale bar, 100 μm. **(D)** Quantification of EGFP mRNA expression 48h after transfection with EGFP constructs together with/without den12a-VPR. Values shown as mean with n = 3.

**Figure S19.**
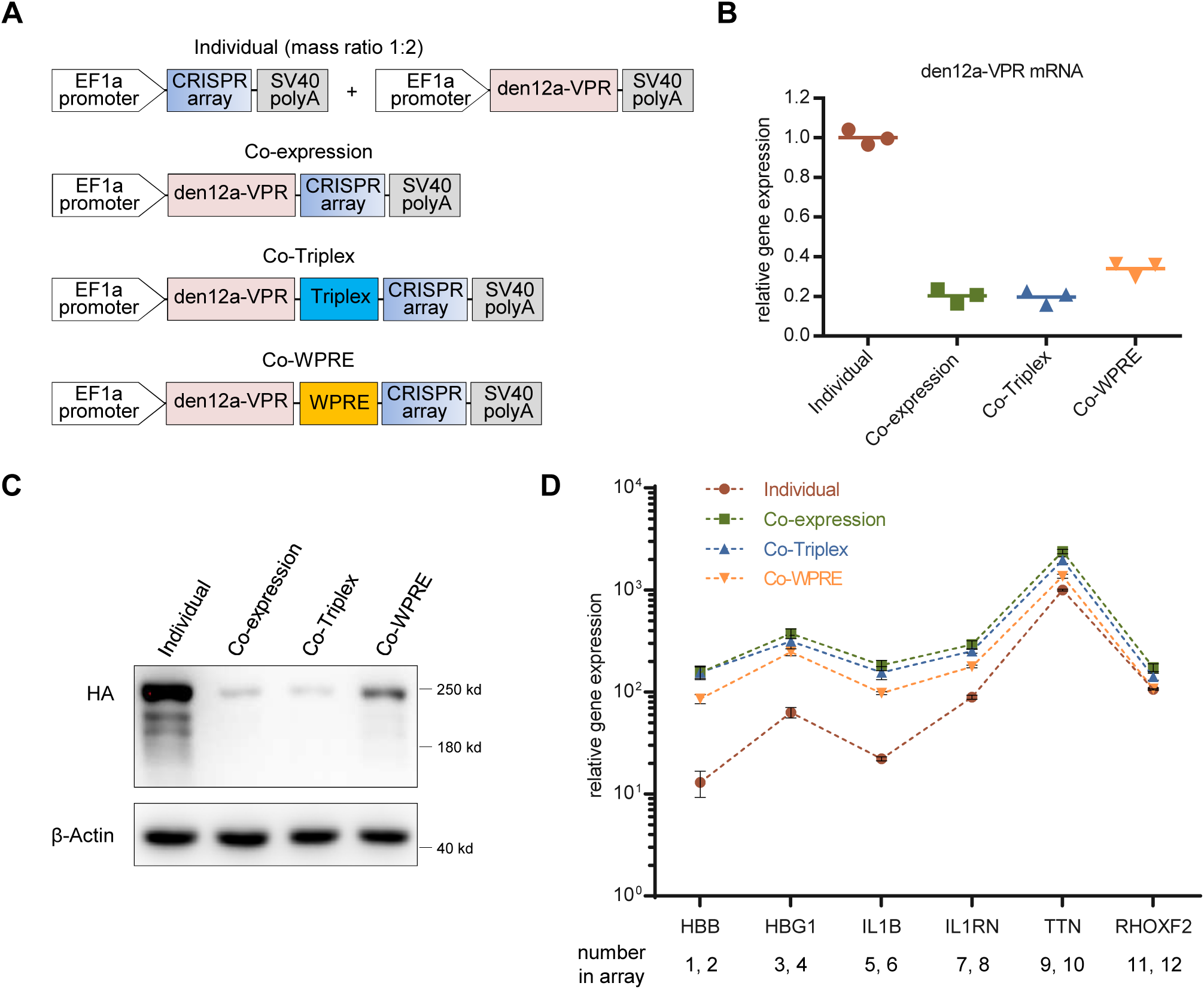
Co-expression of Cas protein and CRISPR array on a single transcript. (**A**) Schematics of constructs used to assess the targeting efficiency of co-expressing Cas protein and CRISPR array on a single transcript. (**B**, **C**) Relative mRNA (**B**) and protein (**C**) expression level of denAsCas12a-VPR in HEK293T cells 48h after transfection with constructs depicted in (**A**). Values shown as mean with n = 3. (**D**) Quantification of relative mRNA expression over non-targeting control in HEK293T cells 48h after transfection with constructs depicted in (**A**). Values shown as mean ± sd with n = 3.

**Figure S20.**
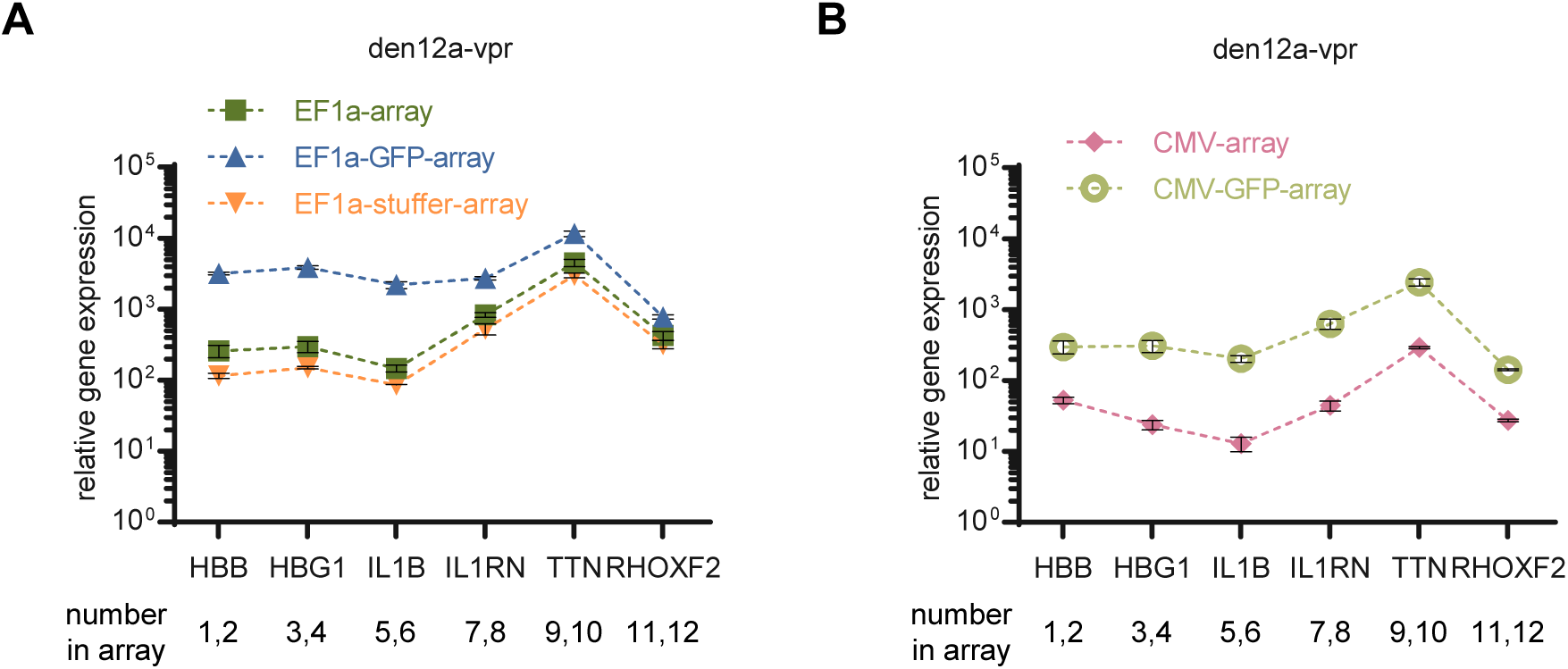
Expression of Pol II promoter-driven CRISPR arrays can be influenced by certain upstream insertions. **(A)** Quantification of relative mRNA expression over non-targeting control in HEK293T cells 48h after transfection with EF1a-den12a-VPR and EF1a-driven arrays with/without upstream EGFP or stuffer. The 745bp stuffer with a GC content of ∼30% is cloned from the pLKO.1 cloning vector (see Supplementary Table for detailed sequence). Values shown as mean ± sd with n = 3. **(B)** Quantification of relative mRNA expression over non-targeting control in HEK293T cells 48h after transfection with EF1a-den12a-VPR and CMV-driven arrays with/without upstream EGFP. Values shown as mean ± sd with n = 3.

**Figure S21.**
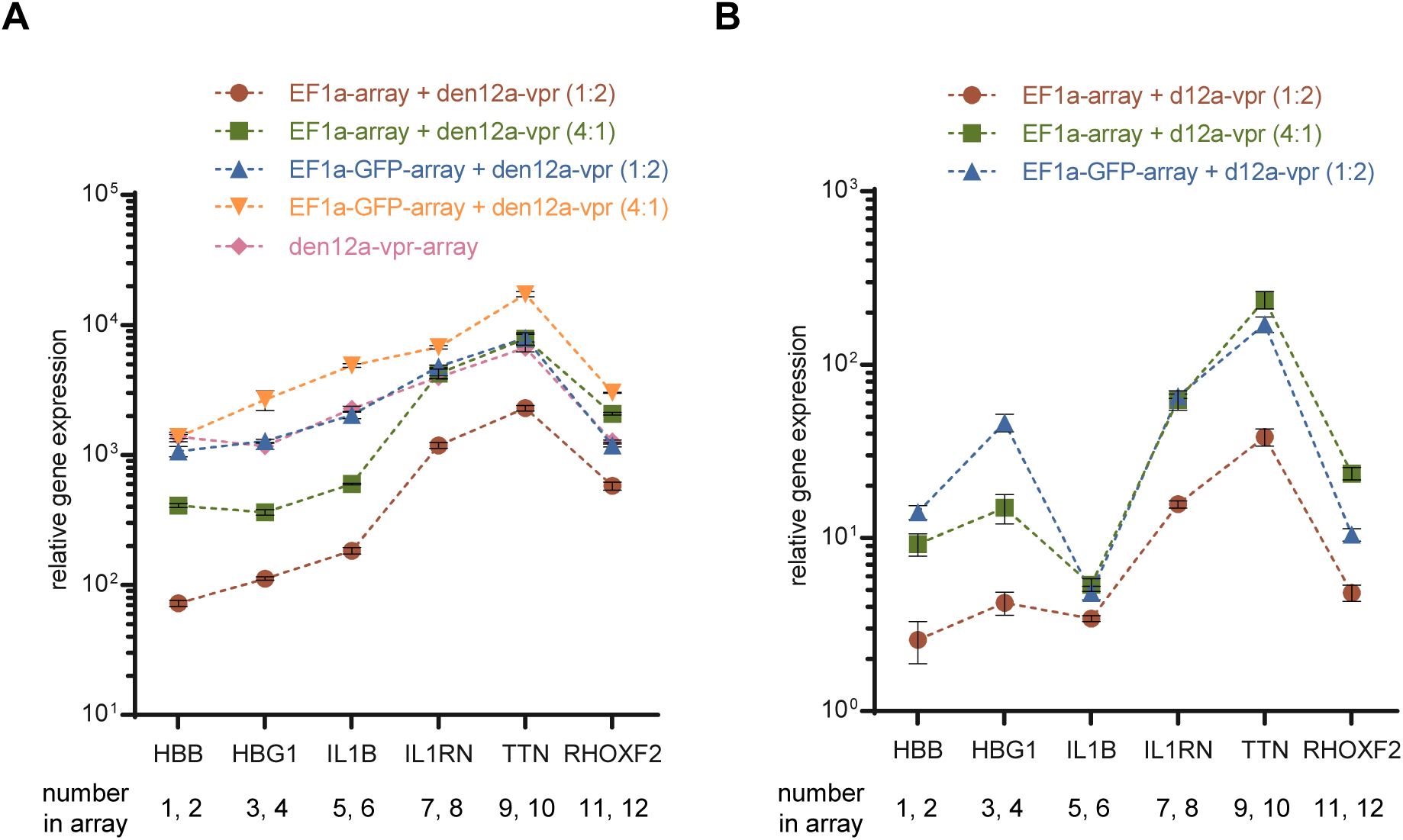
Comparison of the targeting efficiency across different mass ratios or expression patterns. **(A)** Quantification of relative mRNA expression over non-targeting control in HEK293T cells 48h after transfection with the indicated constructs at the mass ratios annotated in parentheses. **(B)** Quantification of relative mRNA expression over non-targeting control in HEK293T cells 48h after transfection with dAsCas12a-VPR and the indicated array-expressing constructs at the mass ratios annotated in parentheses. Values shown as mean ± sd with n = 3.

**Figure S22.**
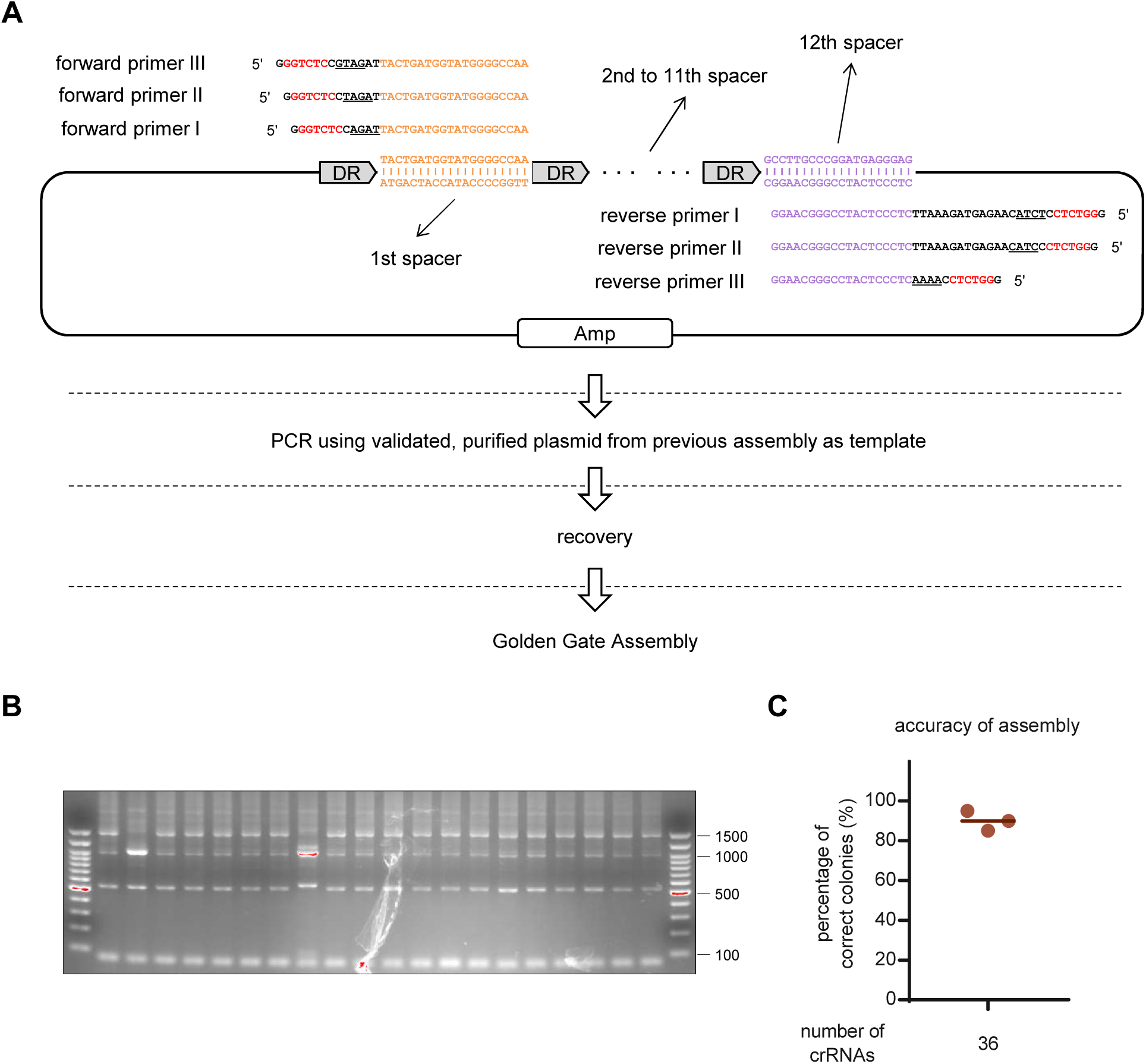
Generation of an array containing 36 crRNAs by two rounds of assembly. **(A)** Schematic illustration and workflow of the second round of assembly for the generation of an array containing 36 crRNAs. A detailed description of the first round of assembly is illustrated in Figure 1 and will not be shown here again. The resulting plasmid from the first round of assembly, encoding an array of 12 crRNAs, is used as a template for the second round of 3 PCR reactions (with primer pair I, II, III, respectively). Recovery the PCR products. Then, set up and run a standard Gold Gate assembly reaction with purified segments and destination cloning vector. **(B)** Representative images of colony PCR to evaluate the accuracy of the second round of assembly, and the corresponding accuracy are shown in **(C)**.

**Figure S23.**
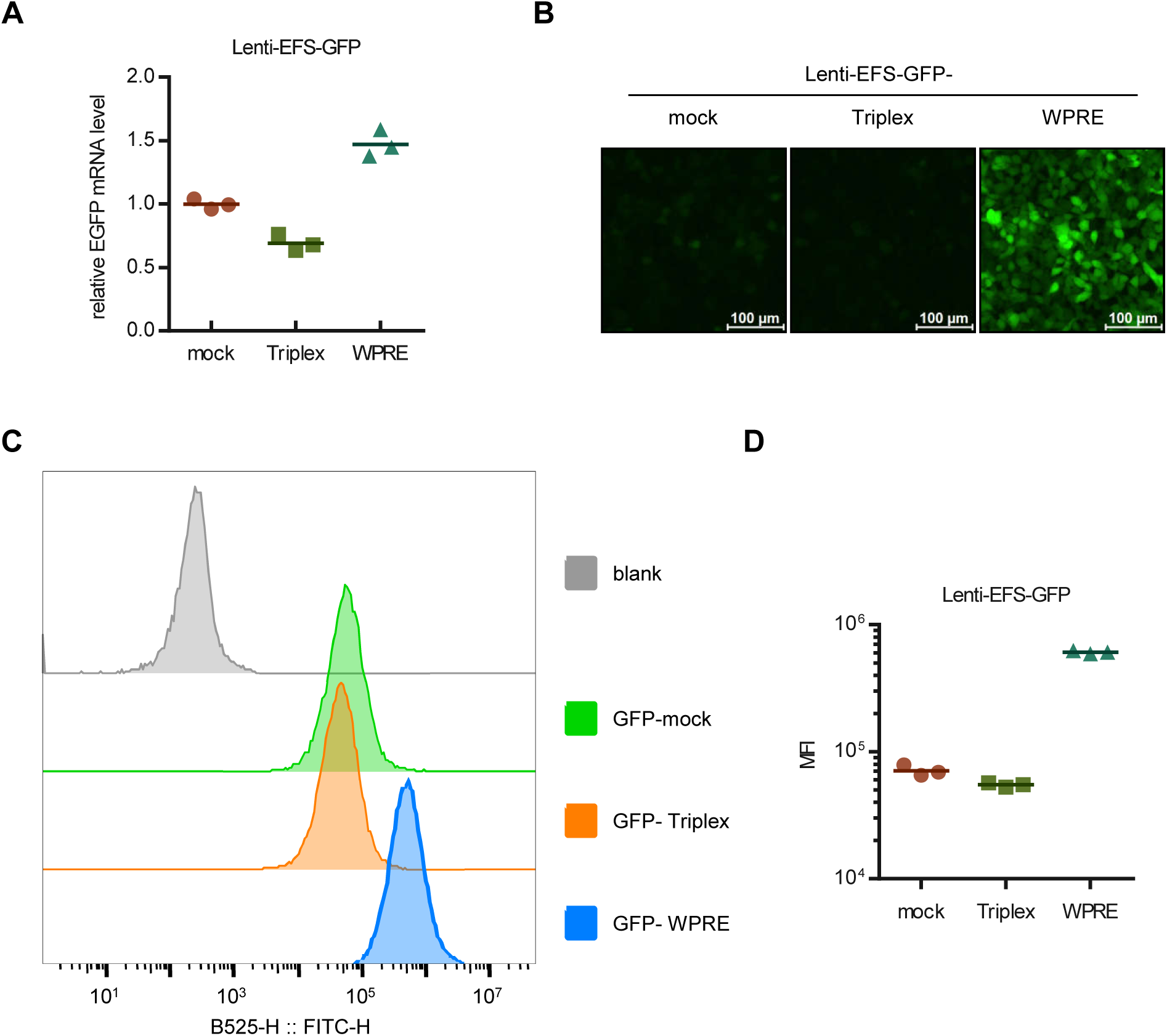
Introduction of WPRE, rather than the “Triplex” element, enhances the expression of gene delivered by lentivirus. **(A-D)** Quantification of EGFP mRNA expression **(A)**, representative fluorescence images **(B)**, representative fluorescence histograms detected by flow cytometry **(C)**, and mean fluorescence intensities **(D)** in HEK293T cells 72h after infection with EGFP-encoding lentivirus, either with or without WPRE or the “Triplex” element. Scale bar, 100 μm. Values shown as mean with n = 3.

## Notes

### Competing Interest Statement

The authors have declared no competing interest.

